# Drivers of Rift Valley fever virus persistence and the impact of control measures in a spatially heterogeneous landscape: the case of the Comoros archipelago, 2004–2015

**DOI:** 10.1101/2021.03.10.434721

**Authors:** Warren S. D. Tennant, Eric Cardinale, Catherine Cêtre-Sossah, Youssouf Moutroifi, Gilles Le Godais, Davide Colombi, Simon E. F. Spencer, Mike J. Tildesley, Matt J. Keeling, Onzade Charafouddine, Vittoria Colizza, W. John Edmunds, Raphaёlle Métras

## Abstract

Rift Valley fever (RVF) is one of the many zoonotic arboviral haemorrhagic fevers present in Africa. The ability of the pathogen to persist in multiple geographically distinct regions has raised concerns about its potential for spread to and persistence within currently disease-free areas. However, the mechanisms for which RVF virus persistence occurs at both local and broader geographical scales have yet to be fully understood and rigorously quantified. Here, we developed a mathematical metapopulation model describing RVF virus transmission in livestock across the four islands of the Comoros archipelago and fitted this model in a Bayesian framework to surveillance data conducted in livestock across those islands between 2004 and 2015. In doing so, we estimated the importance of island-specific environmental factors and animal movements between those islands on the persistence of RVF virus in the archipelago, and we further tested the impact of different control scenarios on reducing disease burden. We demonstrated that the archipelago network was able to sustain viral transmission over 10 years after assuming only one introduction event during early 2007. Movement restrictions were only useful to control the disease in Anjouan and Mayotte, as Grande Comore and Mohéli were able to self-sustain RVF viral persistence, probably due to local environmental conditions that are more favourable for vectors. We also evidenced that repeated outbreaks during 2004-2020 may have gone under-detected by local surveillance in Grande Comore and Mohéli. Strengthened longterm and coordinated surveillance would enable the detection of viral re-emergence and evaluation of different relevant vaccination programmes.

## Introduction

Rift Valley fever (RVF) is a zoonotic arboviral haemorrhagic fever of increasing global health concern. In most cases, it is asymptomatic in humans, but in some cases, it can cause dengue-like symptoms, or in rare instances, more severe conditions such as meningo-encephalitis, haemorrhagic fever or death. In domestic ruminant livestock (cattle, sheep and goats), RVF virus infections cause waves of abortions and high neonatal deaths [1, 2]. RVF was described for the first time in Kenya in 1931 [3]. Since then, the disease has been reported throughout Africa, and outside the African continent in Madagascar (1979), in the Arabian Peninsula (2000) and in the Comoros archipelago (2007) [4–6]. Beyond its potential for spread to further geographical areas, a major concern is the likelihood of persistence in previously disease free regions [7–11]. These persistence mechanisms vary between ecosystems depending on local host communities and meteorological factors allowing favourable conditions for mosquito vectors to complete their life cycle and to be capable of virus transmission [12]. Whilst these mechanisms may apply within a geographically limited homogeneous ecosystem, over a larger geographical scale, one needs to account for spatial heterogeneity. This includes considering other factors and mechanisms such as the variability of the environmental conditions impacting vector transmission, or the movements of hosts across space [13-16].

Previous modelling studies for RVF have focused on estimating key transmission mechanisms in single patch systems, e.g. Mayotte [17, 18], or on viral spatial spread during or between epidemics, e.g. in South Africa and Uganda [19, 20]. However, no study to date has estimated the importance of both environmental variables and animal movements on RVF virus persistence in a spatially heterogeneous system, by fitting a mathematical model to disease data, precluding the formal assessment of disease control measures in a real-world settings. In order to better understand the mechanisms of RVF viral spread and persistence in a spatially heterogeneous system, we developed and fitted a metapopulation model to a series of RVF seroprevalence studies in livestock across the Comoros archipelago—a collection of four islands located in the South-Western Indian Ocean, between Madagascar and Mozambique.

In this paper, we thus sought to (i) estimate the importance of island-specific variables and animal movements across the islands on RVF spread, (ii) assess the likelihood of RVF persistence in the system without re-introduction from mainland Africa or Madagascar, and (iii) assess the impact of livestock movement control measures on disease incidence in the Comoros archipelago. To do this, we developed a mathematical metapopulation model to describe the spread of RVF virus within and between the four islands of the Comoros archipelago (Figure 1), and fitted this model in a Bayesian framework to livestock seroprevalence data collected from 2004 until 2015. Consequently, we estimated the basic reproduction number of the disease on each island over time, and the mean annual number of livestock which move between the four islands. We then used our model to forecast specific RVF antibody prevalence on all four islands in the absence of another explicit introduction event of the virus from outside the Comoros archipelago. Finally, we assessed the impacts of movement restrictions and reducing within-island transmission on each island upon the total number of new infections in livestock from 2004 to 2015.

**Figure 1:**
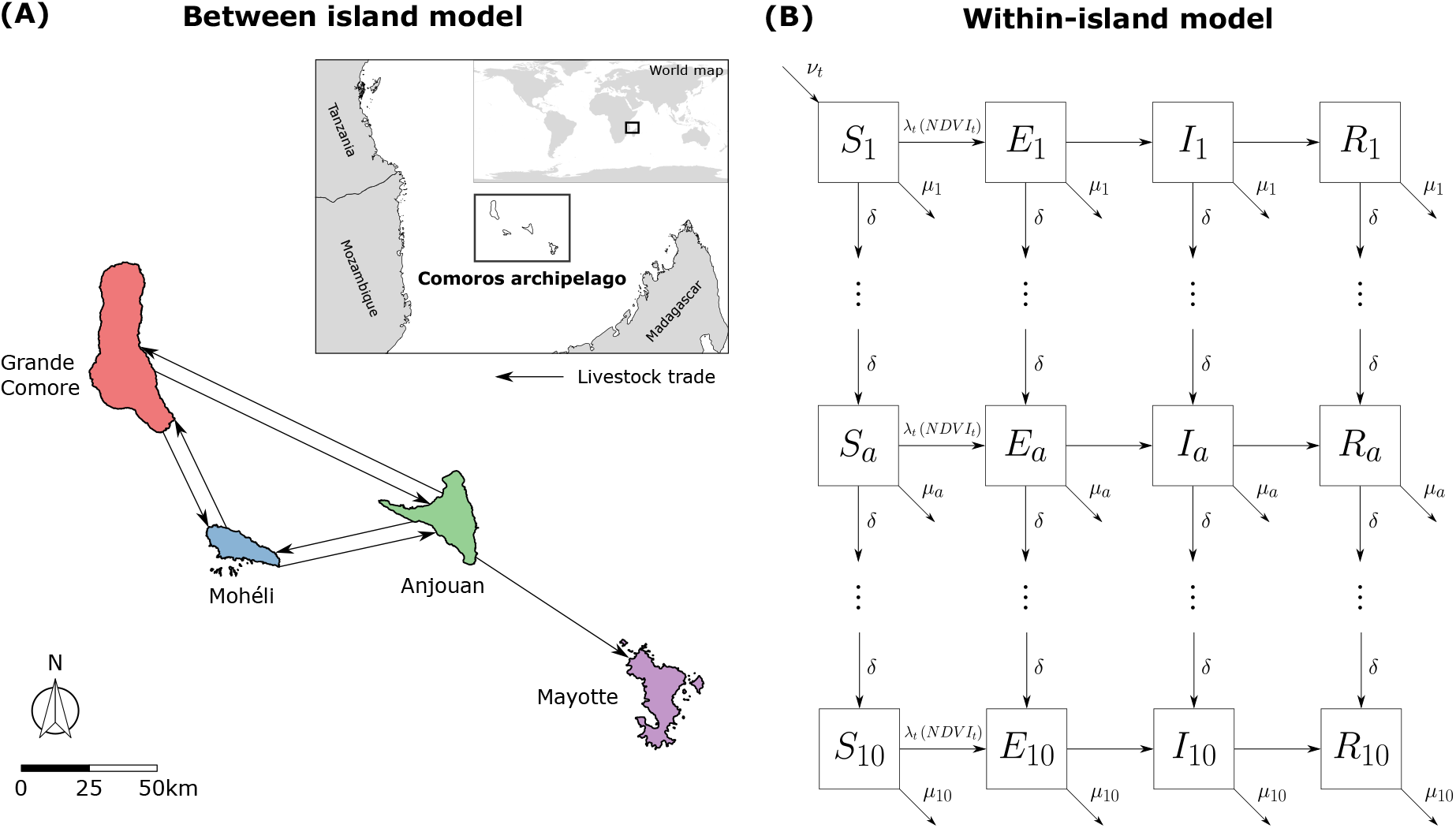
Metapopulation model for RVF virus transmission in the Comoros archipelago. In order to quantify the drivers of Rift Valley fever in the Comoros archipelago, we developed a metapopulation model describing RVF virus infection of livestock (cattle, sheep, goats) in the Comoros archipelago. We modelled **(A)** the explicit movement of livestock (solid black arrows) between the four islands in the Comoros archipelago: Grande Comore, Mohéli, Anjouan and Mayotte. **(B)** Within-island viral transmission was modelled as an age-stratified Susceptible-Exposed-Infected-Recovered (SEIR) model, including demographics (birth, ageing, and deaths) and transmission driven by Normalized Difference Vegetation Index (NDVI). For further details and notation, refer to Methods & materials.

## Results

### Livestock seroprevalence data

We used age-stratified RVF IgG seroprevalence data collected in livestock as part of several sero-surveys conducted amongst the four islands of the archipelago (namely Grande Comore, Mohéli, Anjouan and Mayotte). A total of 8,423 samples—2,191 in Grande Comore, 475 in Mohéli, 857 in Anjouan and 4,900 in Mayotte—were collected over a 12 year period (July 2004-June 2015). Summary statistics for these data are shown in Figure 2. For details on these data, refer to Methods & Materials.

**Figure 2:**
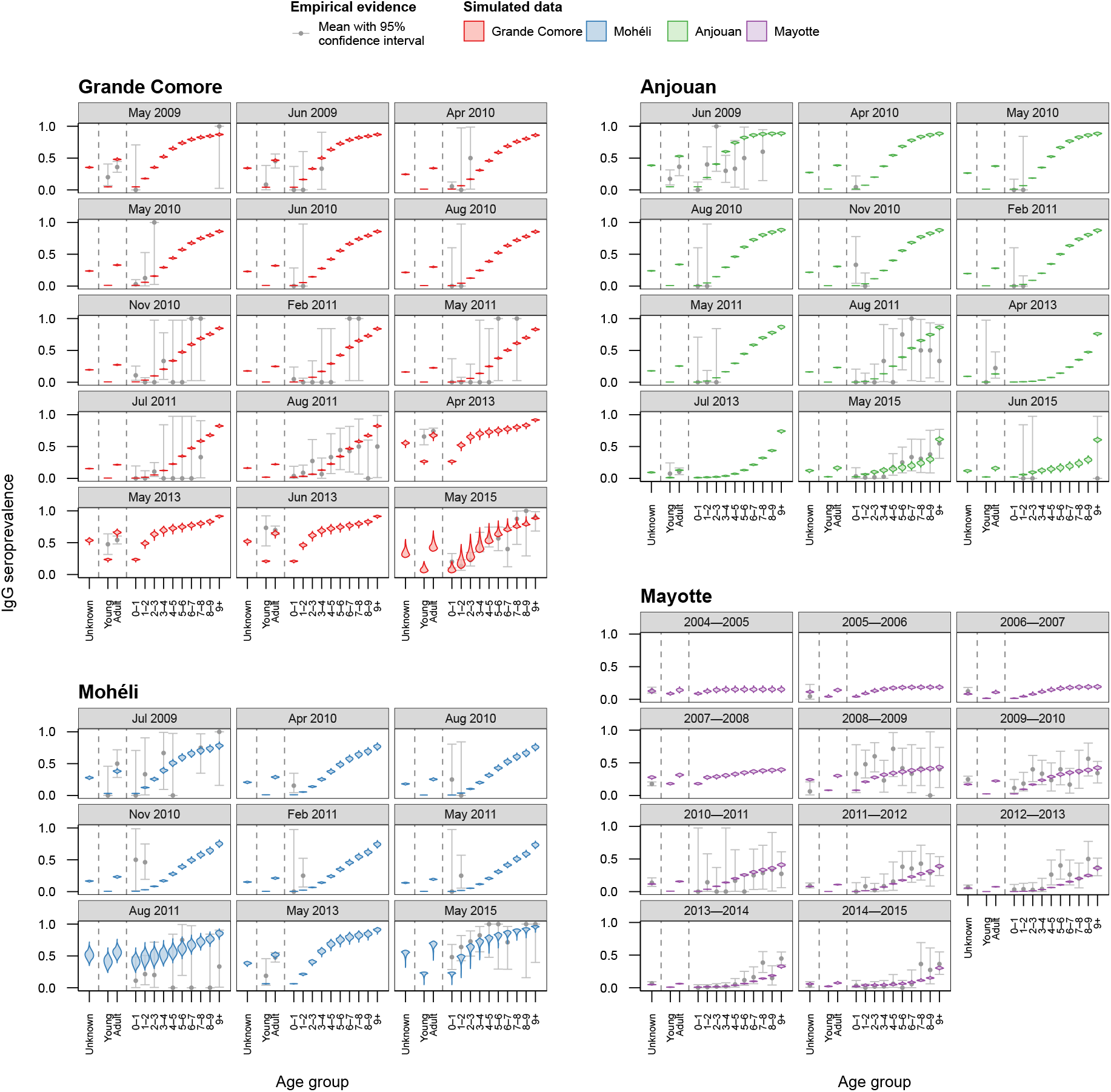
Model fit of the best fitted model (Model 3b) for each sero-survey conducted between July 2004 and June 2015. The exponential transmission model with the same seasonal component *α* and different baseline transmission *β* for each island fitted to the data best out of all five models tested (DIC = 1,189). Shown is the fitted simulated IgG seroprevalence (coloured violins) for each aggregated sero-survey conducted throughout the study period. Seroprevalence of Grande Comore (red), Mohéli (blue) and Anjouan (green) were aggregated by month, and seroprevalence for Mayotte (purple) was aggregated by year. The grey dots show the observed age-stratified IgG seroprevalence with 95% confidence interval (vertical bars). Simulated seroprevalence was generated through 1,000 realisations of the metapopulation model.

### Estimation of island-specific transmission and animal movements

We modelled RVF viral transmission within age-structured livestock populations within each island as a function of the Normalized Difference Vegetation Index (NDVI), and between islands through the movement of livestock. Five models with different relationships between NDVI and within-island transmission were fitted in a Bayesian framework to the age-stratified livestock seroprevalence data in order to estimate island-specific viral transmission rates. We used Deviance Information Criterion (DIC) [21] to discriminate the relative quality of fitted models (Supplementary Table 1). We present the fit for the tested models in Figure 2 and Supplementary Figures 1-4 — showing the comparison between simulated age-stratified IgG seroprevalence in livestock against the available serological data.

The model assuming a similar exponential relationship between NDVI and transmission amongst livestock across islands, with island-specific baseline transmission values (Model 3b) fitted to the empirical data with the greatest accuracy according to DIC (Figure 2). Predictions of Model 3b included the observed rise in seroprevalence of RVF in Grande Comore and Mohéli and fall in seroprevalence in Anjouan between 2011 and 2014. Furthermore, in the complete absence of age information (Mayotte 2004-2008) the model captured the rise in seroprevalence in 2007-2008. There were only a few serological surveys for which the model was unable to capture. These discrepancies included sero-surveys in young livestock conducted in Grande Comore from April until June 2013.

The full set of parameter estimates for Model 3b can be found in Supplementary Table 2 and Supplementary Figure 5. These parameter estimates corresponded to a (median) maximum annual seasonal reproduction number, *R_st_*, on each island from highest to lowest was 3.99 for Grande Comore (95% credible interval (CrI) = [3.17, 4.72]), 3.40 for Mohéli (95% CrI = [2.33, 5.54]), 2.95 for Anjouan (95% CrI = [2.15, 3.83]) and 2.77 for Mayotte (95% CrI = [2.36, 3.14]). The geometric mean of seasonal reproduction numbers was also greater than one across all four islands (Supplementary Table 3). Consequently, our model inferred multiple outbreaks to have occurred on both Grande Comore and Mohéeli: Mohéli in 2011, Grande Comore in 2012 and both Mohéli and Grande Comore in 2014 (Supplementary Figure 6). In addition, the importation of infectious animals was inferred to begin between December 2006 and April 2007 (with 95% credibility) with 4.15 (95% CrI = [1.33, 7.10]) infectious animals being introduced each week. Livestock were also traded between islands within the Comoros archipelago (Figure 3). Mayotte was estimated to be the largest importer of animals, with 1,875 (95% CrI = [1,707, 2,058]) importations per annum, all of which came from Anjouan. As a consequence, Anjouan was the largest exporter, exporting 2,912 (95% CrI = [2,728, 3,104]) animals per year, with 629 (95% CrI = [565, 689]) and 408 (95% CrI = [365, 449]) to Grande Comore and Mohéli respectively. Mohéli exported and imported similar number of animals per year: 739 (95% CrI = [685, 795]) imports and 719 (95% CrI = [641, 800]) exports. The majority of Mohéli’s exports were trade to Grande Comore, 657 (95% CrI = [588, 726]), with approximately half that imported, 331 (95% CrI = [294, 367]).

**Figure 3:**
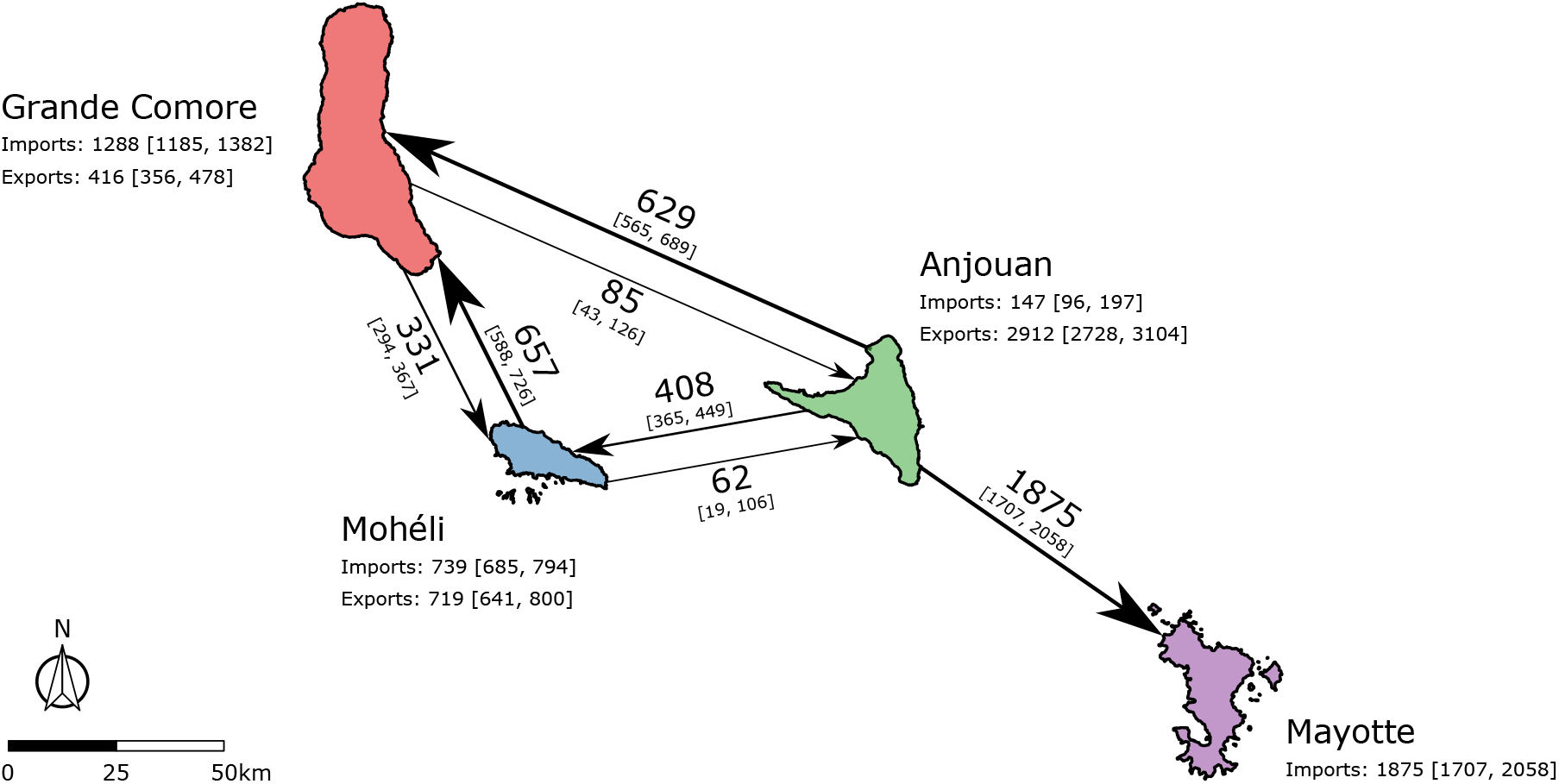
Estimated livestock trade network in the Comoros archipelago from best fitted model (Model 3b), presented as the annual number of livestock heads moved between islands. The imports and exports of each island in the Comoros archipelago were estimated by fitting the metapopulation model to age-stratified sero-surveys conduct from July 2004 until June 2015. The estimated annual trade network of livestock in the Comoros archipelago is shown, with the direction of each arrow indicating the direction of trade between islands. The median and 95% credible interval (CrI) of estimated annual livestock movements are shown on each arrow.

### RVF transmission dynamics and persistence in the Comoros archipelago

Based on 1,000 realisations of the best model (Model 3b), we observed a rise in seroprevalence during 2007-2008 on all four islands (Figure 4). This was due to high seasonal reproduction numbers on each island attributed to high NDVI values (Supplementary Figure 7) occurring alongside a sufficiently high proportion of susceptible animals; or related to the importation of infectious livestock into Grande Comore from the African mainland during late 2006 and early 2007, following the RVF outbreak in East Africa [22].

**Figure 4:**
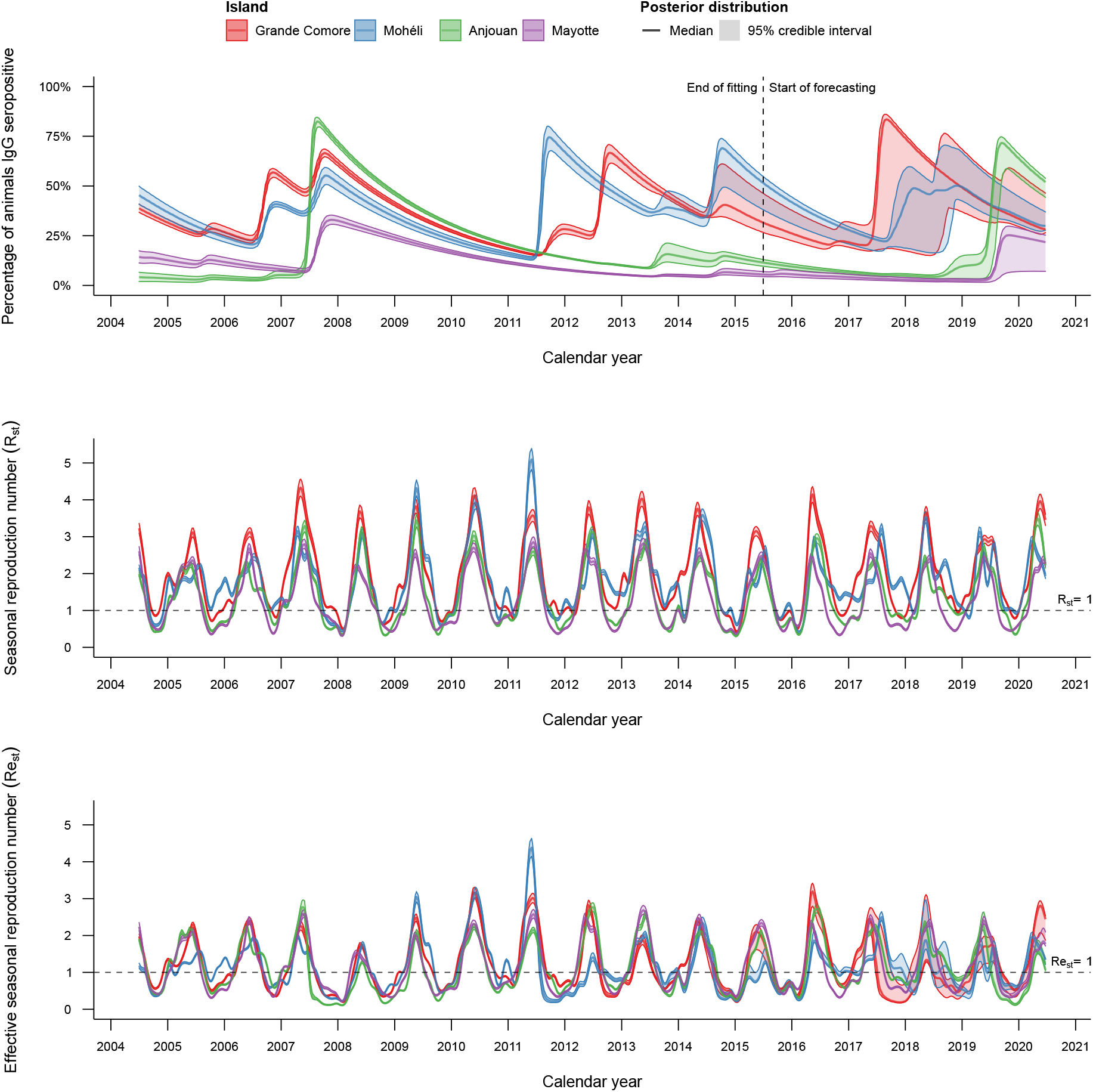
Simulated RVF IgG seroprevalence on each island in the Comoros archipelago, 2004 202(), for the best fitted model (Model 3b). The metapopulation model was fitted to age-stratified serosurveys from July 2004 until June 2015. Fitted models were then simulated until July 2020. Shown is the median and 95% credible interval of **(A)** IgG seroprevalence, **(B)** the seasonal reproduction number (*R_st_*) and **(C)** the effective seasonal reproduction number (*Re_st_*) for Grande Comore (red), Mohéli (blue), Anjouan (green) and Mayotte (purple). Distributions shown were estimated from 1,000 realisations of the metapopulation model.

Forecasting beyond July 2015 until June 2020 showed outbreaks occurring on all four islands again, but their timing and magnitude varied greatly between islands. According to our model predictions, large outbreaks of RVF occurred on Grande Comore and Mohéli in either 2017 and/or 2018. A small outbreak was predicted for Anjouan in 2018, followed by a substantial epidemic in Anjouan and Mayotte during 2019. By the end of June 2020, the model predicted population level seroprevalence to be 28.3% on Grande Comore (95% CrI = [27.0%, 43.1%]), 30.2% on Mohéli (95% CrI = [27.7%, 36.6%]), 51.4% on Anjouan (95% CrI = [44.0%, 53.9%]) and 20.7% on Mayotte (95% CrI = [7.5%, 26.0%]), giving weight to the hypothesis that the Comoros archipelago is able to sustain RVF viral transmission without an explicit introduction of the virus from mainland Africa or Madagascar.

### Impact of livestock movement control measures

To further investigate the role of animal movements on the epidemiology of Rift Valley fever in the Comoros archipelago, we compared the total number of livestock infections in the 2004–2015 period under different movement restriction scenarios (Figure 5A). Under the full trade network, the estimated number of infections per island were 362,659 (95% CrI = [341,277, 420,691]) in Grande Comore, 57,244 (95% CrI = [50,664, 60,630]) in Mohéli, 98,567 (95% CrI = [91,241, 104,117]) in Anjouan and 7,482 (95% CrI = [6,532, 8,527]) in Mayotte.

**Figure 5:**
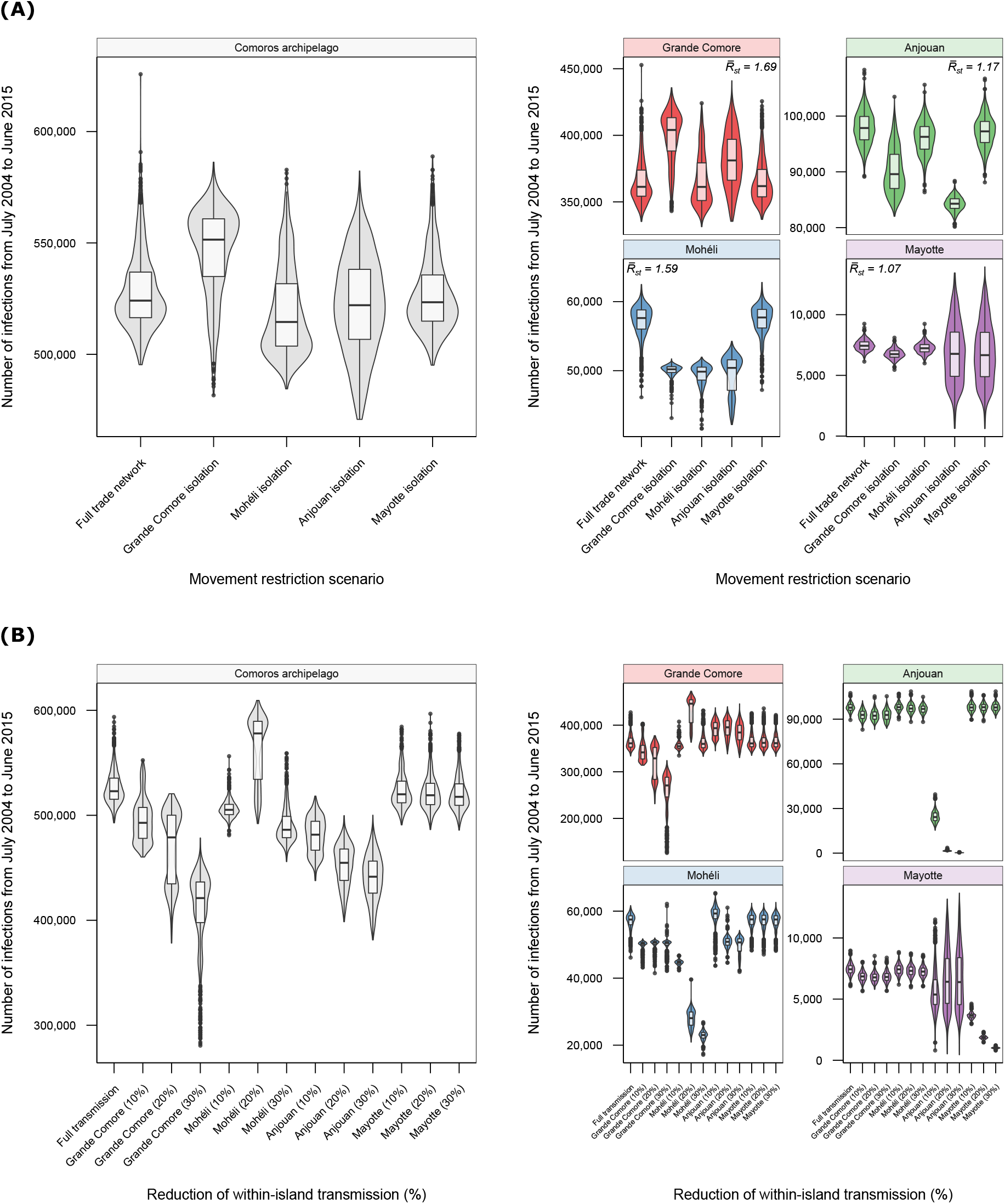
Effect of control measures on the number of infections in the Comoros archipelago from 2004—2015. The total number of infections from July 2004 until June 2015 is shown under each control measure scenario on each island in the Comoros archipelago (grey): Grande Comore (red), Mohéli (blue), Anjouan (green) and Mayotte (purple). **(A)** Full (100%) restrictions on imports and exports were placed on each island. For each island, the median estimate of the geometric mean seasonal reproduction number, 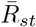, independent of movement restrictions is shown. **(B)** Within-island transmission rate was reduced by 10%, 20% and 30% on each island independently. Distributions were generated from 1,000 realisations of each scenario.

#### Grande Comore

Movement reductions on Grande Comore generated a median increase of 20,076 in the total number of cases across all four islands compared to the full trade network. Relative to the (median) total number of infections on each island under the full trade network, reducing the number of imports and exports into Grande Comore by 100% increased the number of infections in Grande Comore itself by 10% (95% CrI = [-4.0%, 18.5%]). This increase was due to a delayed 2012-2013 outbreak in Grande Comore, resulting in a small (new) outbreak in 2013–2014 and more severe outbreak in 2014–2015 on the island (Supplementary Figure 8). Under 100% reduction in Grande Comore’s exports, the total number of infections in Mohéli reduced by 12.6% (95% CrI = [10.6%, 18.8%]). Furthermore, the total number of infections in Anjouan and Mayotte decreased by 10% on average under full movement restrictions on Grande Comore.

#### Mohéli

Isolating Mohéli from Grande Comore and Anjouan decreased the (median) total number of infections across all four islands by 15,150. Compared to the full trade network, Mohéli’s own total number of infections fell by 14% (95% CrI = [10.0%, 25.8%]). The total number of infections on the other three islands—Grande Comore, Anjouan and Mayotte—were unaffected.

#### Anjouan

The total number of infections in the Comoros archipelago reduced by 1,998 on average compared to the full trade network under complete movement restrictions to and from Anjouan. Restricting imports and exports of Anjouan by 100%, only reduced its own total number of infections by 14.1% (95% CrI = [11.6%, 17.3%]), and reduced the total number of infections on Mohéli by approximately 12.2%.

#### Mayotte

Restricting movement from Anjouan to Mayotte increased the total number of cases across all four islands on average: a rise of 724 infections over the study period. A complete reduction in imports into Mayotte averted the 2007 Mayotte epidemic, but may have instead caused an outbreak in 2011 owing to sufficient local conditions for transmission (Supplementary Figure 8). As a result of the movement restrictions to Mayotte, infections on Grande Comore and Mohéli (the first and third most populous islands) increased by 0.5% on average.

### Effects of reducing within-island transmission

To investigate the long-term impacts of island-specific control measures, such as vector control, on the dynamics of RVF throughout the Comoros archipelago, we compared the total number of livestock infections in 2004-2015 period under 10%, 20% and 30% reductions in the transmission rate of each island (Figure 5B).

#### Grande Comore

Reducing the within-island transmission rate on Grande Comore by 10%, 20% and 30% caused a median decrease in the total number of cases across all four islands by 31,000, 44,882 and 102,693 compared to the full transmission model. These overall decreases in incidence was because of a reduction in the number of cases on Grande Comore itself and Mohéli: decreases of 25% (95% CrI = [15.4, 56.4]) and 12.2% (95% CrI = [10.5%, 20.7%]) on Grande Comore and Mohéli respectively under 30% control. In each scenario, the 2006 and 2012 simulated outbreaks on Grande Comore were not as severe and the 2011 outbreak on Mohéli was delayed until 2013 (Supplementary Figure 9).

#### Mohéli

Both a 10% and 30% reduction in the transmission rate on Mohéli reduced the number of cases throughout the Comoros archipelago by 18,626 and 37,632 on average over the study period respectively. However, under a 20% reduction in the transmission rate, the number of cases increased by 54,154 on average. This is because under the 20% scenario, susceptibility is high enough (higher than the full transmission model) for a more severe outbreak to occur on Mohéli in mid-2011 despite the reduction in transmission intensity. This larger outbreak on Mohéli resulted in a larger outbreak in Grande Comore through trade of infected livestock (Supplementary Figure 10). As a consequence, the total number of cases on Grande Comore increased by 23.5% (95% CrI = [3.5, 28.7]) compared with the full transmission model under 20% control on Mohéli.

#### Anjouan

The total number of infections in the Comoros archipelago was reduced by 42,280, 69,114 and 82,344 from July 2004 until June 2015 under 10%, 20% and 30% reductions in the transmission rate on Anjouan respectively. Almost all of these reductions occurred on Anjouan: a 98.5% (95% CrI = [97.8, 99.1%]) decrease in the number of cases on Anjouan under 30% control levels. Similar to the movement restriction scenario on Anjouan, the 2007 Mayotte epidemic did not occur under each control scenario (Supplementary Figure 11).

#### Mayotte

Under 10%, 20% and 30% transmission reduction scenarios, the number of cases on Mayotte decreased by 51.1% (95% CrI = [44.3%, 57.3%]), 75.0% (95% CrI = [71.5%, 78.3%]) and 86.5% (95% CrI = [84.7%, 88.3%]) respectively. The temporal epidemiological dynamics of the other three islands were unaffected under control on Mayotte (Supplementary Figure 12).

## Discussion

Understanding the transmission dynamics of Rift Valley fever (RVF) within animal populations is essential towards estimating human spillover risk and assessing the impact of control measures [23, 24]. Characterising persistence mechanisms for RVF are useful to assist long-term surveillance programmes, anticipate re-emergence and assess the impact of control measures [25]. In spatially heterogeneous systems, these persistence mechanisms not only include factors at a local scale, but also those over a larger geographical scale, such as pathogen reintroduction from neighbouring regions [26]. However, the effects of hosts, vectors and their environment on the persistence of RVF are not yet well understood or quantified. In order to improve understanding of how environmental factors and animal trade influence the transmission dynamics of Rift Valley fever, we developed a metapopulation model for RVF infection in livestock and fitted it to data in a multi-insular ecosystem — the Comoros archipelago — from serological data during 2004-2015.

We showed that (i) the within-island RVF virus transmission was similarly driven by NDVI across these islands, (ii) the virus was able to persist across the network over 12 year period, (iii) within-island controls were more effective than livestock movement restrictions between islands. Finally, we provided evidence that some recent outbreaks may have gone undetected.

In total, five different metapopulation models were fitted: each with their own model describing the rate of transmission between livestock. The transmission mechanisms mapping our seasonal driver, Normalized Difference Vegetation Index (NDVI), to transmission rate in an exponential manner fitted to the data best according to Deviance Information Criterion (DIC). This result not only agrees with an NDVI-driven transmission model for RVF applied to Mayotte only [17], but suggests that the effects of our choice of covariate (NDVI), alongside island-specific baseline transmission rates, were sufficient to explain the observed serology on each island. The estimated baseline transmission rates on each island from highest to lowest were Grande Comore, Mohéli, Anjouan and Mayotte. This variation may be attributed to the different agricultural ecosystems on each island, such as livestock production system or within-island trade network of varying intensity [27–29]. These findings were reflected in the (time-varying) reproduction number: a metric to quantify disease severity and threshold criterion to determine endemicity [30].

Across all four islands, the geometric mean seasonal reproduction number was greater than one, indicating persistence of RVF virus in the Comoros archipelago. The maximum estimated reproduction number for each island was between 2.5 and 4, which is in line with previous reproduction number estimates for RVF [20, 31, 32]. The seasonal reproduction number was greater than one for over three quarters of the year on Grande Comore and Mohéli, and only half the year on Anjouan and Mayotte (Supplementary Table 3), reflecting island-specific conditions. Lower reproduction number estimates on Anjouan and Mayotte might explain why outbreaks were less likely to occur on these islands, as time periods with conditions suitable for RVF transmission (high NDVI) were offset by time periods with poor transmission suitability (low NDVI). This explanation at least agrees with existing evidence that an explicit introduction event from outside the Comoros archipelago may have been essential for an outbreak to occur on Mayotte in 2006/2007 [5, 6, 17].

Importation of RVF infected (or RVF IgM antibody positive) animals from outside the Comoros archipelago may have also been an essential factor in causing the 2018 epidemic in Mayotte [18, 33]. In the absence of an explicit re-introduction event of the RVF virus, our model predicted that an outbreak was ongoing in Mayotte one year after the serological data suggests. Indeed, sequence analysis of the strains confirmed that the strain circulating in Mayotte from 2018 onwards was that responsible for the RVF epidemics in Uganda in 2017 [34]. Yet even without an explicit re-introduction of the virus into Grande Comore within our framework, our model predictions suggest that the environmental conditions of Grande Comore and Mohéli were sufficient to cause substantial RVF outbreaks in 2017 and 2018. There is little empirical evidence to support these predictions as it stands, owing to the lack of recent active surveillance on these islands, suggesting these outbreaks may have been missed. As a consequence, any vaccination campaign that would be reactive to the detection of outbreaks would currently be ineffective unless surveillance was first improved.

Our study showed that restricting imports into Mayotte prevented an epidemic in 2007 (Supplementary Figure 8). This suggests that at least for Mayotte, the importation of livestock from Anjouan to Mayotte is paramount for inducing large outbreaks on the island. Indeed, viral re-emergence on Mayotte resulted from viral re-introductions (likely from infected animals imported from neighbouring islands), coupled with an important proportion of the local livestock being susceptible to infection [17, 18, 35]. However, in contrast to previous data that showed Mayotte by itself in a closed ecosystem (without animal imports) could not sustain viral transmission between its two epidemics [8, 17], our results indicated that an outbreak may have occurred during 2011 on Mayotte instead due to sufficient environmental conditions for transmission.

We demonstrated that island-specific control strategies, such as movement restrictions or vector control, may result in more poor epidemiological outcomes over the long term. For example, restricting livestock movements to and from Grande Comore only served to delay an outbreak to a season which was more suitable for transmission, resulting in a more severe outbreak. As Grande Comore was also the island least affected by the control measures of the other three islands, our results may suggest that only within-island control (perhaps with some combination of movement restrictions) ought to be considered to reduce disease burden on Grande Comore. However, Mohéli’s temporal dynamics were found to be very sensitive to controls implemented on Grande Comore or Anjouan. Island-specific controls on Mayotte affected the other three islands the least, and thus only Mayotte is appropriate for focused island-specific control measures. A combination of the controls may be more appropriate to implement on Grande Comore, Mohéli and Anjouan simultaneously, but the potential impacts on the complete system (including Mayotte) should be thoroughly assessed first.

There are some limitations to using a mechanistic approach to elucidate the factors that drive spread and persistence of RVF. Our findings of RVF being endemic in the Comoros archipelago may be partially attributed to our choice of modelling framework: a deterministic framework in which disease can technically never be eradicated. However, rather than discussing persistence of RVF as an absence of stochastic extinction, we have presented our claims of persistence in terms of the reproduction number. That is, our results show that, *on average*, RVF has the ability to remain endemic on the Comoros archipelago provided that the climate conditions remain favourable to support transmission. We also did not explicitly model mosquitoes within our framework as there were insufficient data on all competent vectors of RVF in the Comoros archipelago, or elsewhere, to parametrise such an approach [27, 36, 37]. It is clear however that understanding mosquito population dynamics and their vectorial capacity for RVF virus will be paramount to fully disentangle the seasonal drivers of RVF from one another. Our model can be further extended to include the population dynamics of RVF vectors in areas where such data are available.

In summary, we have presented the first metapopulation model for RVF fitted to empirical data. We have shown that the virus is able to persist in the Comoros archipelago without the need for the explicit introduction of the virus from eastern Africa or Madagascar. Moreover, we have identified that the ecological features of Grande Comore and Mohéli were more suitable to maintain the virus, whereas livestock trade and low seroprevalence levels were essential for RVF epidemics to occur on Anjouan and Mayotte. Our results suggest that several outbreaks occurred in Grande Comore and Mohéli that were missed owing to insufficient surveillance. This study emphasises the importance of sustaining long-term, coordinated surveillance programmes in order to elicit an early enough vaccination control response to avert epidemics in livestock and resultant spill-over into human populations.

## Methods & materials

### Study area

The Comoros archipelago is a group of four islands (Grande Comore (1,146 km^2^), Mohéli (290 km^2^), Anjouan (424 km^2^) and Mayotte (374 km^2^)) located in the northern part of the Mozambique Channel, between Mozambique and Madagascar, populated with about 1 million inhabitants [29, 38]. The climate of the islands is marine tropical, and the Comoros archipelago are old volcanic islands, with varying ecosystems [29, 39, 40]. The livestock (cattle, sheep and goat) production system is extensive, and the total animal population is estimated to be over 350,000 [41]. Animals are raised for local consumption. Grande Comore, Mohéli and Anjouan are part of the Union of the Comoros, and may exchange animals regularly. Mayotte has been a French department since 2011 and a EU outermost territory since 2014, and no official import of animals are reported from the neighbouring island, Anjouan, whilst some unreported imports occur on a regular basis [12, 27, 39, 40].

### Livestock seroprevalence data

We analysed cross-sectional seroprevalence data conducted in the Union of Comoros as part of surveillance programs in Grande Comore, Mohéli and Anjouan as presented by Roger et al. [42] and Roger et al. [27]. Surveys were conducted in 2009, 2012, 2013 and 2015. The livestock prevalence data from Mayotte covered the period 2004 to 2016 and resulted from the SESAM (Système d’épidémio-surveillance animale à Mayotte) surveillance system as presented in Métras et al. [12] and Métras et al. [17].

### Metapopulation model

To decipher the relative roles of meteorological factors and livestock movements on RVF persistence across the four islands of the Comoros archipelago, we developed a metapopulation model, using an island as a patch. Within each of the four patches, we modelled RVF transmission dynamics and islands were connected by animal movements. A schematic summarising the main components of the model is shown in Figure 1.

The model was deterministic and discrete-time, where each time step was one week. The livestock populations (cattle, sheep and goats) were modelled for each island in the Comoros archipelago, with each split into 10 age groups *a*: 0-1 (*a* = 1), 1-2 (*a* = 2),…, and 9+ (*a* = 10) years old. Animals died with an age-dependent rate *μ_a_*, and each island *i* contained *N_i,a_* animals of each age group. Animals of each age group *a* moved from island *i* to island *j* at a weekly rate *m_t,ij,a_*, where *m_t,ii,a_* denoted the number of animals of age group *a* which remained on island *i* at time *t*. Animals were born into the youngest age group at rate *ν_t,i_* on each island *i* and aged at rate *δ*. For simplicity, we assumed that the total livestock population of each island was constant over time.

Animals of each age group *a* and island *i* were classified as either susceptible (*S_i,a_*), exposed (*E_i,a_*), infectious (*I_i,a_*) or recovered (*R_i,a_*) to the virus. We assumed that animals were exposed to the virus for one week, and remained infectious for one week. Once recovered, animals were immune to infection for the duration of their life. We assumed no maternal protection to the virus, and thus new animals were assumed to be fully susceptible to the disease. At each time *t*, susceptible animals became infected at an island-dependent rate *λ_t,i_*. Infectious animals of age *a* were introduced into each island *i* from sources outside the Comoros archipelago at a time-dependent rate denoted by 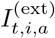.

Given livestock population sizes at time *t*, the total number of susceptible, exposed, infectious and recovered livestock for each age group *a* and island was calculated at subsequent time t + 1 as follows:

For animals less than one year old (the first age group, *a* = 1):

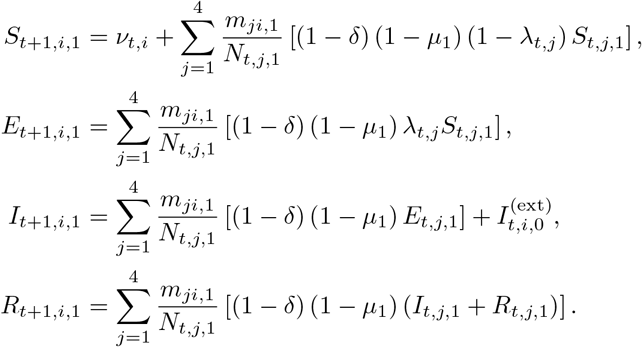

For animals between one and nine years old (age groups *a* = 2,…, 9):

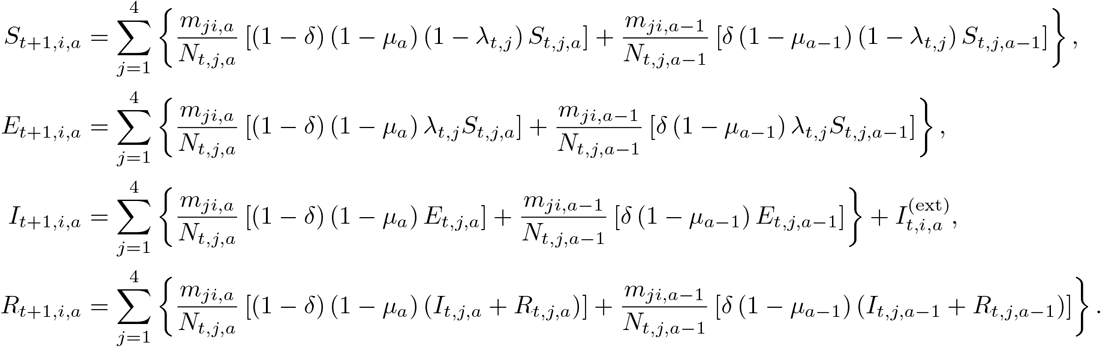

For animals greater than nine years old (the final age group, *a* = 10):

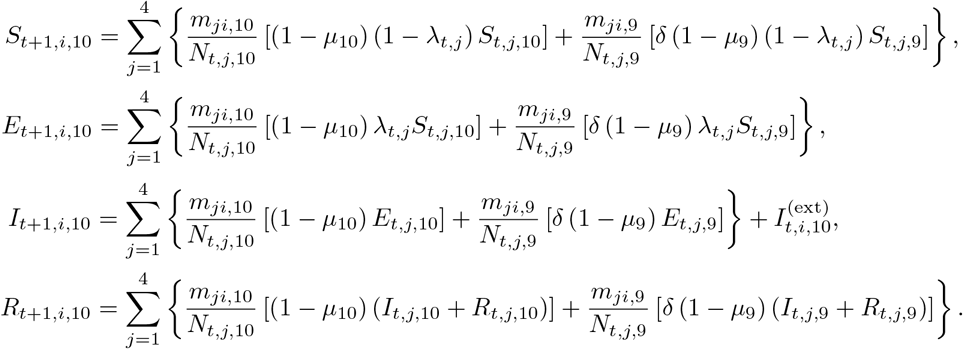

The birth rate, *ν_t,i_*, and force of infection, *λ_t,i_*, were defined as:

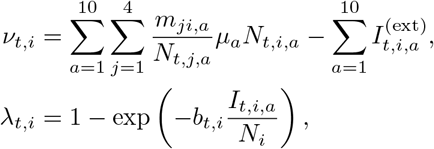

for any island *i* and time-dependent transmission rate *b_t,i_*.

At the first time point (*t* = 0), a small number of exposed and infectious animals were initialised on each island. A proportion of the total population on each island, *ϵ_i_*, was also assumed to be immune at the start of the simulation. It was also assumed that the age of immune animals was proportional to the age of the population. Therefore, for each island *i*,

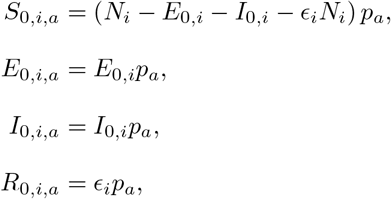

where *p_a_* denotes the proportion of the livestock population in age-group *a*.

### Movement between islands

We assumed that the total number of movements between islands was constant over time. Owing to the substantial distances between each island, we assumed that animals could move along the network motivated by Roger et al. [27]. The movement network is shown in Figure 1. That is, only the following movements were possible: Grande Comore to Mohéli, Grande Comore to Anjouan, Mohéli to Grande Comore, Mohéli to Anjouan, Anjouan to Grande Comore, Anjouan to Mohéli, and Anjouan to Mayotte. All animals that did not move on this network were assumed to remain on their respective islands.

As the size of adult livestock relative to the size of boats used to travel between islands was large, only the two youngest age groups were moved between islands. We also assumed that the number of movements per age group *a* was proportional to the total number of animals in each age group at time *t* and the total number of weekly movements between island *i* and island *j*.

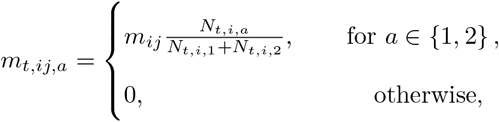

where *m_ij_* denoted the total number of weekly livestock movements between island *i* and island *j*.

### External importation of infection

On account of the RVF outbreak in eastern Africa between 2006 and 2007, we assumed that infectious animals were imported into Grande Comore for a limited time. As with movements between islands in the Comoros archipelago, we assumed that only the two youngest age groups were imported. We also assumed that the age of imported animals was proportional to the age of the Comoros archipelago livestock population. Given the weekly number of infectious animals imported, 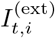, the number of infectious imports per island *i* and age group *a* was defined as follows:

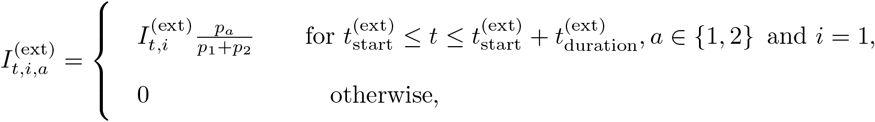

where 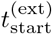 and 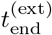 denote the start and end of the importation window respectively.

### Seasonally driven transmission functions

We used Normalized Difference Vegetation Index (NDVI) as a proxy for the effects of seasonal mosquito population dynamics on RVF transmission rates. The estimated NDVI at each time point and island was obtained by aggregating NDVI estimates of 250m by 250m grid squares for each island [43]. Weekly estimates were obtained by smoothing the aggregated estimates with a Gaussian kernel of 21 days [44]. As the relationship between NDVI and transmission rate is unknown, we tested three underlying models for transmission: linear, exponential and constant as presented below.

#### Linear

The transmission rate scaled linearly with NDVI:

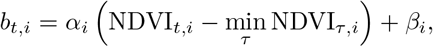

for some positive parameter *α_i_* and *β_i_* representing the baseline level of transmission on each island.

#### Exponential

The transmission rate scaled exponentially with NDVI:

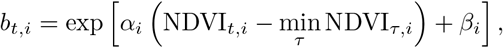

for some positive parameter *α_i_* and exp (*β_i_*) denoting the baseline level of transmission on each island.

#### Constant

As a baseline comparison, we also modelled a time-independent transmission rate, thus not depending on NDVI:

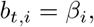

where *β_i_* denotes the island dependent transmission rate.

As the exposure and infectious periods were fixed at one week, the seasonal reproduction number and effective seasonal reproduction numbers for each island were given by *R_st,i_*, = *b_t,i_* and 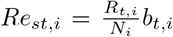 respectively. An island *i* was deemed to allow the virus to persist if the geometric mean of *R_st,i_* over the time period was greater than 1.

#### Parameters

The parameters of the model were selected based on the current knowledge on RVF epidemiology and demography of the livestock population in the Comoros archipelago. The parameters that were fixed in the model are shown in Table 1. To allow for monthly aggregates of RVF IgG seroprevalence to be calculated (used in model fitting), the time steps of the model represent 1.08 calendar weeks. Therefore, the weekly ageing rate δ was chosen such that animals took 48 time steps (one year) to age. The mortality rates of age groups 1-9 and 10 were set to 8.8 × 10^-3^ and 6.2 × 10^-3^ respectively [45, 46]. We assumed that mortality rates of livestock were independent of the island on which they were kept. The population size of each island was calculated by aggregating estimates of sheep, goat and cattle population estimates for each island using the Gridded Livestock Map of the World [41]. All other parameters were estimated by fitting the model to serological surveys carried out on each island from July 2004 until June 2015.

**Table 1:**
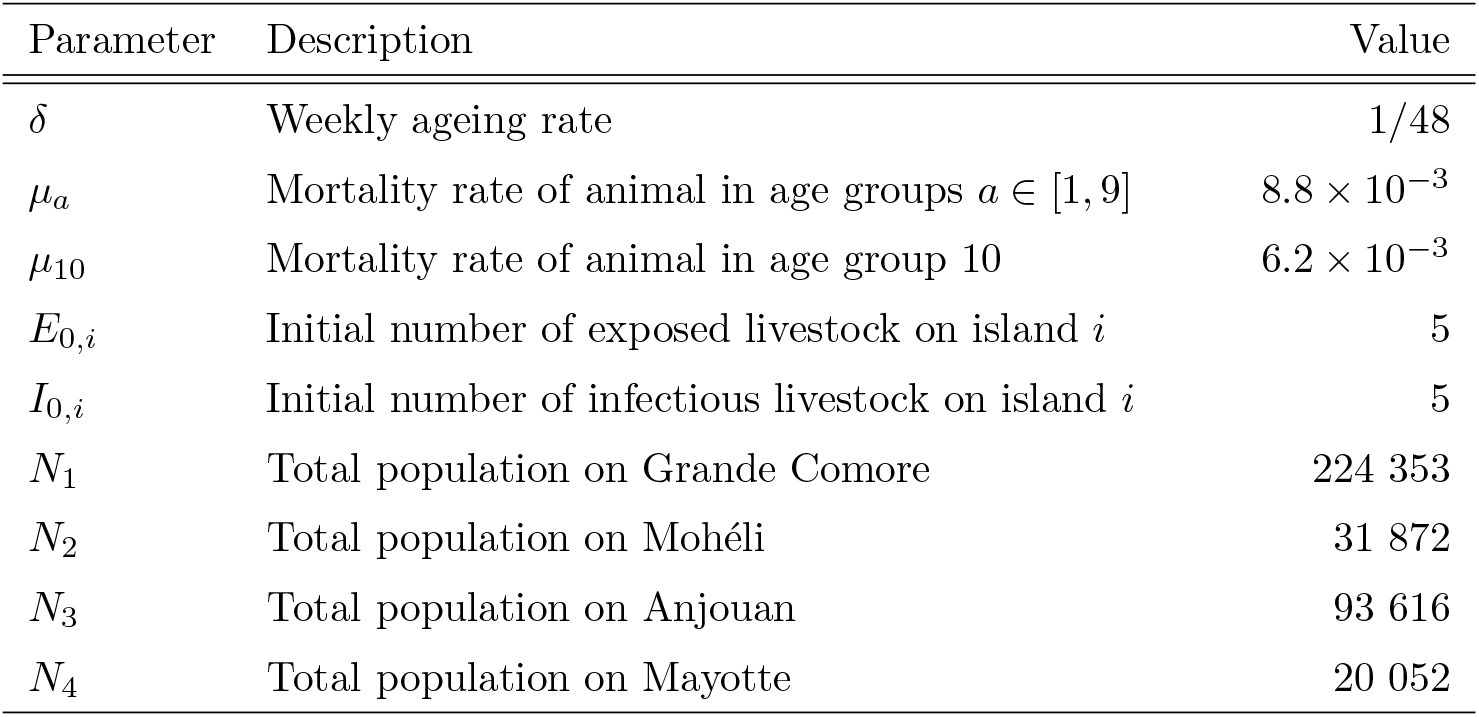
Fixed model parameters. The parameters in the metapopulation model were chosen based on the current understanding of the epidemiology of Rift Valley fever and the demography of the livestock population in the Comoros archipelago. The remaining parameters were estimated by fitting the model to the serological data collected between 2004 and 2015.

| Parameter | Description | Value |
| --- | --- | --- |
| $\delta$ | Weekly ageing rate | 1/48 |
| $\mu_a$ | Mortality rate of animal in age groups $a \in [1, 9]$ | $8.8 \times 10^{-3}$ |
| $\mu_{10}$ | Mortality rate of animal in age group 10 | $6.2 \times 10^{-3}$ |
| $E_{0,i}$ | Initial number of exposed livestock on island $i$ | 5 |
| $I_{0,i}$ | Initial number of infectious livestock on island $i$ | 5 |
| $N_1$ | Total population on Grande Comore | 224 353 |
| $N_2$ | Total population on Mohéli | 31 872 |
| $N_3$ | Total population on Anjouan | 93 616 |
| $N_4$ | Total population on Mayotte | 20 052 |

### Model fitting and parameter estimation

To estimate the remaining parameters *θ* of the model, we fitted the metapopulation model to the serological data in a Bayesian framework.

As farms and animals were randomly sampled for serological testing on each island, we assumed that the number of RVF specific IgG antibody positive animals was binomially distributed given the total number of animals tested and the proportion of the livestock population that were immune at the time of testing. For Grande Comore, Mohéli and Anjouan, surveys were aggregated by the month in which they were conducted. As surveys in Mayotte were conducted throughout the year, we aggregated surveys by epidemiological year, which began in July. Using the meta-population model to estimate the proportion of the population that were immune at the age of testing,

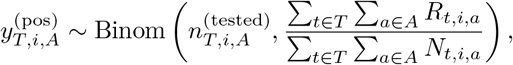

where 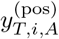 and 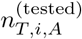 denote the number of animals that were bled and tested RVF antibody positive in age group(s) *A* on island *i* over the aggregated time window *T*, respectively. For surveys that were not age-specific, animals were classed as either adults (age groups 2-10) and infants (age group 1). In surveys where the age of animals was unknown, all age groups in the model were used to calculate the proportion immune at the time of testing. As each survey was conducted independently from one another, the likelihood was given by the product of observing each survey.

The priors for each parameter estimated are shown in Supplementary Table 4. Normal priors were chosen for the movement parameters, with prior parameters selected based on consultation with the Comorian veterinary services and previously reported inter-island trade estimates [17, 27]. The priors for the proportion of the livestock population immune on each island was set to be a beta distribution with mean seroprevalence of 10% on Anjouan and Mayotte, and 40% on Grande Comore and Mohéli [42]. The variance of each beta distribution was determined after consultation with local experts. Weak normal priors were placed on transmission parameters as these are yet to be robustly quantified. Normal priors were used for the start and duration of the import into Grande Comore, with prior parameters selected to correspond to reports of the RVF outbreak in Kenya during 2006/7 [47, 48]. A weak normal prior was used for the size of the import as there is scarce information on infectivity of imported animals during 2006/2007.

We sampled from the posterior distribution of the parameters *θ*, using an adaptive Markov Chain Monte Carlo Metropolis-Hastings random walk algorithm [49, 50]. Parameters were estimated for five epidemiological models:

1. Model 1: constant transmission with different *β_i_* for each island,
2. Model 2a: linear transmission with different *α_i_* and the same *β* for each island,
3. Model 2b: linear transmission with the same *α* and different *β_i_* for each island,
4. Model 3a: exponential transmission with different *α_i_* and the same *β* for each island, and
5. Model 3b: exponential transmission with the same *α* and different *β_i_* for each island.

Convergence was assessed through visual inspection of the trace plots and calculation of the Gelman-Rubin 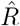 statistic [51]. The best model was determined by which one had the lowest deviance information criterion (DIC) [21].

### Forecasting and control scenarios

To assess the ability for RVF to persist within the Comoros archipelago beyond 2015, we forecast RVF virus seroprevalence in livestock using empirical NDVI data from 2015 until 2020. We did this by drawing 1000 samples from the joint posterior distribution of the best model fit and using these parameters to simulate the model until July 2020. To assess the importance of inter-island trade on the transmission of RVF within the archipelago, we imposed (100%) movement restrictions (imports and exports) on each island. To investigate the potential long-term impacts of island-specific control measures, such as vector control, on the transmission of RVF within the archipelago, we reduced the transmission rate by 10%, 20% and 30% on each island independently. For each control scenario, we calculated the total number of infections in livestock on each island compared with the total number of infections on the full movement and transmission network.

## Acknowledgements

We thank Matthieu Roger, Floriane Boucher, the veterinary services and the surveyors for their help in collecting livestock trade data in the Union of the Comoros. We thank the veterinary services of Mayotte for sharing their surveillance data. This study was partly supported by the TROI project (FEDER Interreg V) under the DP One health Indian Ocean (www.onehealth-oi.org) in partnership. The study was conducted under the Vaccine Efficacy Evaluation for Priority Emerging Diseases (VEEPED) project. WSDT, WJE, MJK, MJT and SEFS are funded by the Department of Health and Social Care using UK Aid funding managed by the National Institute for Health Research (Vaccine Efficacy Evaluation for Priority Emerging Diseases: PR-OD-1017-20007). The views expressed in this publication are those of the authors and not necessarily those of the Department of Health and Social Care.

