## Supplementary information for "Drivers of Rift Valley fever virus persistence and the impact of control measures in a spatially heterogeneous landscape: the case of the Comoros archipelago, 2004–2015"

### Supplementary Tables

**Supplementary Table 1: Model selection.** Five models were fitted to the age-stratified sero-surveys from July 2004 to June 2015. The deviance information criterion (DIC) is presented for each model. The model with the lowest DIC was determined to be the best fitted model (indicated by *). The table shows the model identified,underlying assumptions on viral transmission, and whether the seasonal and constant transmission components were the same or different for each island.

| Model number | Transmission | Seasonal component for each island ($\alpha$) | Constant component for each island ($\beta$) | DIC |
|---|---|---|---|---|
| 1 | Constant | - | Different | 1,580 |
| 2a | Linear | Different | Same | 1,433 |
| 2b | Linear | Same | Different | 1,578 |
| 3a | Exponential | Different | Same | 1,386 |
| 3b* | Exponential | Same | Different | 1,189 |

**Supplementary Table 2: Summary of estimated parameters for the exponential model (Model 3b).** Parameters of the metapopulation model were estimated by fitting the model to serological surveys conducted on each island through the 2004–2015 period. Shown in the table are the median and 95% credible intervals (CrI) of 10,000 posterior samples for each estimated parameter for the best fitted model (Model 3b).

| Parameter | Description | Median estimate [95% CrI] |
|---|---|---|
| $48m_{12}$ | Annual movement from Grande Comore to Mohéli | 331.0 [293.6, 367.2] |
| $48m_{13}$ | Annual movement from Grande Comore to Anjouan | 84.6 [43.0, 126.2] |
| $48m_{21}$ | Annual movement from Mohéli to Grande Comore | 657.3 [588.0, 726.4] |
| $48m_{23}$ | Annual movement from Mohéli to Anjouan | 62.4 [19.0, 106.5] |
| $48m_{31}$ | Annual movement from Anjouan to Grande Comore | 628.8 [565.1, 689.2] |
| $48m_{32}$ | Annual movement from Anjouan to Mohéli | 408.0 [364.8, 448.6] |
| $48m_{34}$ | Annual movement from Anjouan to Mayotte | 1875.4 [1706.9, 2057.5] |
| $\epsilon_1$ | Proportion immune at time $t = 0$ in Grande Comore | 0.39 [0.37, 0.41] |
| $\epsilon_2$ | Proportion immune at time $t = 0$ in Mohéli | 0.45 [0.40, 0.50] |
| $\epsilon_3$ | Proportion immune at time $t = 0$ in Anjouan | 0.04 [0.02, 0.07] |
| $\epsilon_4$ | Proportion immune at time $t = 0$ in Mayotte | 0.14 [0.11, 0.17] |
| $a$ | Seasonal transmission component scalar for all islands | 8.5 [8.1, 8.9] |
| $b_1$ | Transmission constant for Grande Comore | -0.85 [-0.91, -0.80] |
| $b_2$ | Transmission constant for Mohéli | -0.89 [-0.96, -0.82] |
| $b_3$ | Transmission constant for Anjouan | -1.18 [-1.25, -1.12] |
| $b_4$ | Transmission constant for Mayotte | -1.14 [-1.21, -1.08] |
| $t^{(\text{imp})}_{\text{start}}$ | Start of imports into Grande Comore (epi. weeks since July 2004) | 122 [117, 132] |
| $t^{(\text{imp})}_{\text{duration}}$ | Duration of imports into Grande Comore | 23.6 [10.7, 36.2] |
| $48I^{(\text{imp})}$ | Annual infectious imports into Grande Comore | 199.2 [63.0, 341.0] |

**Supplementary Table 3: Summary statistics of the seasonal reproduction number.** The seasonal reproduction number, $R_{st}$, for RVF based on our fitted model was calculated for each island. The table shows the median and 95% credible interval for the geometric mean seasonal reproduction number over the study period, minimum and maximum annual $R_{st}$ and the number of (calendar) weeks per year for which $R_{st} > 1$ on each island. Estimates were calculated from 1,000 simulations of the best fitted model (Model 3b).

| Island | Mean seasonal reproduction number $R_{st}$ | Minimum annual $R_{st}$ | Maximum annual $R_{st}$ | Number of calendar weeks $R_{st} > 1$ |
|---|---|---|---|---|
| Grande Comore | 1.69 [1.66, 1.72] | 0.87 [0.46, 1.12] | 3.99 [3.17, 4.72] | 39.1 [38.9, 39.8] |
| Mohéli | 1.59 [1.54, 1.64] | 0.87 [0.44, 1.38] | 3.40 [2.33, 5.54] | 40.4 [38.8, 41.4] |
| Anjouan | 1.17 [1.15, 1.19] | 0.48 [0.33, 0.77] | 2.95 [2.15, 3.83] | 28.1 [27.5, 28.5] |
| Mayotte | 1.07 [1.04, 1.10] | 0.48 [0.34, 0.70] | 2.77 [2.36, 3.14] | 25.9 [25.2, 26.3] |

**Supplementary Table 4: Prior distributions for estimated parameters.** The parameters estimated by fitting the metapopulation model to serological surveys from 2004–2015. Priors were chosen based on consultation with the Comorian veterinary services and current and historical understanding of RVF in the Comoros.

| Parameter | Description | Prior |
|---|---|---|
| $48m_{12}$ | Annual movement from Grande Comore to Mohéli | $\text{Normal}(50, 50)$ |
| $48m_{13}$ | Annual movement from Grande Comore to Anjouan | $\text{Normal}\left(350, \frac{350}{10}\right)$ |
| $48m_{21}$ | Annual movement from Mohéli to Grande Comore | $\text{Normal}\left(676, \frac{676}{10}\right)$ |
| $48m_{23}$ | Annual movement from Mohéli to Anjouan | $\text{Normal}(50, 50)$ |
| $48m_{31}$ | Annual movement from Anjouan to Grande Comore | $\text{Normal}\left(625, \frac{625}{10}\right)$ |
| $48m_{32}$ | Annual movement from Anjouan to Mohéli | $\text{Normal}\left(409, \frac{409}{10}\right)$ |
| $48m_{34}$ | Annual movement from Anjouan to Mayotte | $\text{Normal}\left(1500, \frac{1500}{10}\right)$ |
| $\epsilon_1$ | Proportion immune at time $t = 0$ in Grande Comore | $\text{Beta}(20, 30)$ |
| $\epsilon_2$ | Proportion immune at time $t = 0$ in Mohéli | $\text{Beta}(20, 30)$ |
| $\epsilon_3$ | Proportion immune at time $t = 0$ in Anjouan | $\text{Beta}(5, 45)$ |
| $\epsilon_4$ | Proportion immune at time $t = 0$ in Mayotte | $\text{Beta}(5, 45)$ |
| $b_i^{(\text{const})}$ | Transmission constant for island $i$ (constant) | $\text{Normal}(1, 2)$ |
| $a_i^{(\text{linear})}$ | NDVI scalar for island $i$ (linear) | $\text{Normal}(3, 2)$ |
| $b_i^{(\text{linear})}$ | Transmission constant for island $i$ (linear) | $\text{Normal}(1, 2)$ |
| $a_i^{(\text{exp})}$ | NDVI scalar for island $i$ (exponential) | $\text{Normal}(3, 2)$ |
| $b_i^{(\text{exp})}$ | Transmission constant for island $i$ (exponential) | $\text{Normal}(-2, 2)$ |
| $t_{\text{start}}^{(\text{imp})}$ | Start of imports into Grande Comore | $\text{Normal}(128, 24)$ |
| $t_{\text{duration}}^{(\text{imp})}$ | Duration of imports into Grande Comore | $\text{Normal}(24, 12)$ |
| $48I^{(\text{imp})}$ | Annual infectious imports into Grande Comore | $\text{Normal}(150, 150)$ |

### Supplementary Figures

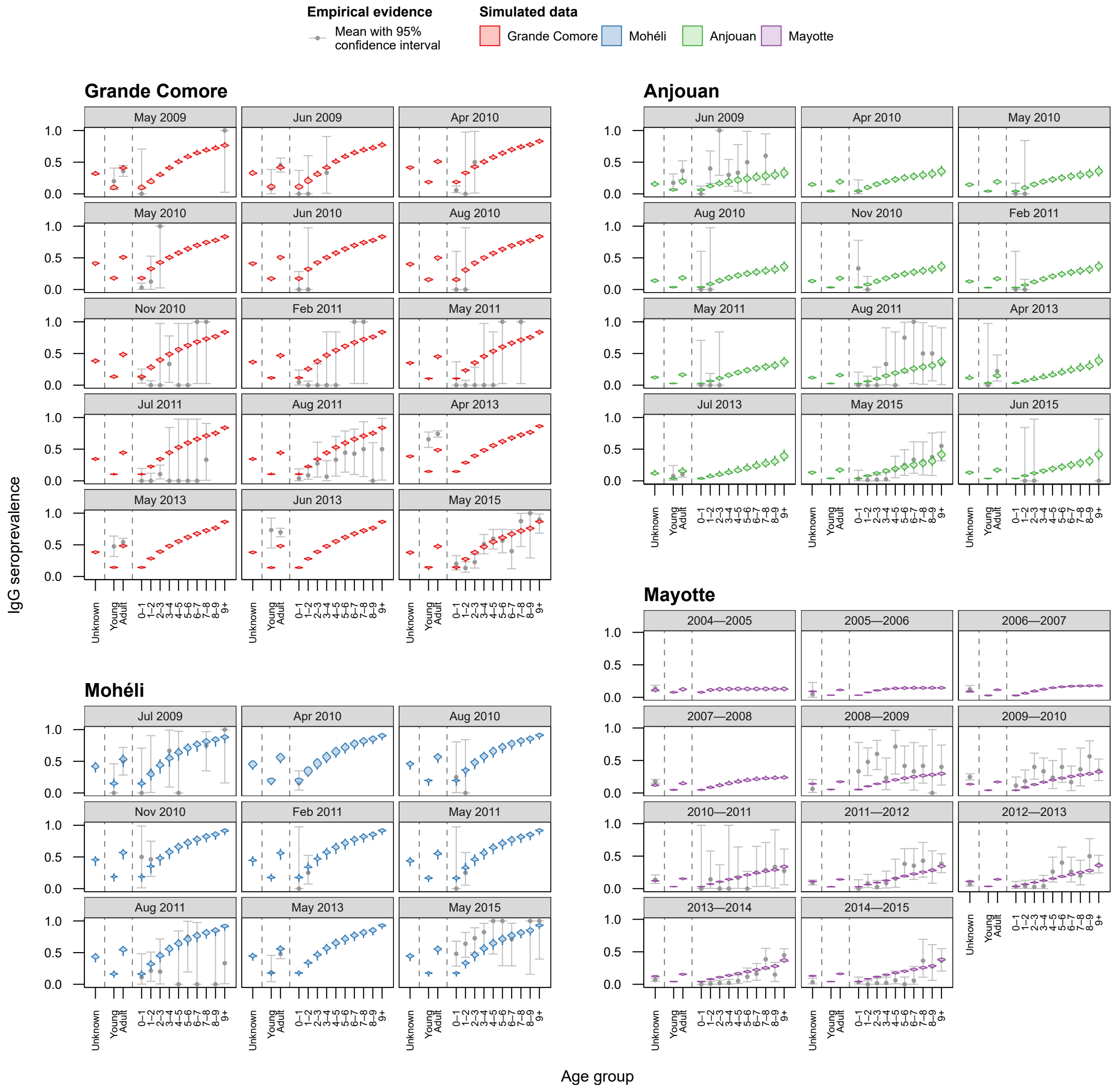


**Supplementary Figure 1: Model fit of the constant transmission model (Model 1) for each sero-survey conducted between July 2004 and June 2015.** The constant transmission model with the different constant transmission $\beta$ for each island fitted to the data worst out of all five models tested (DIC $= 1,580$). Shown is the fitted simulated seroprevalence for each aggregated sero-survey conducted throughout the study period (coloured violins). Seroprevalence of Grande Comore (red), Mohéli (blue) and Anjouan (green) were aggregated by month, and seroprevalence for Mayotte (purple) was aggregated by year. The black dots show the observed age-stratified IgG seroprevalence with 95% confidence interval (vertical bars). Simulated seroprevalence was generated through 1,000 realisations of the metapopulation model.

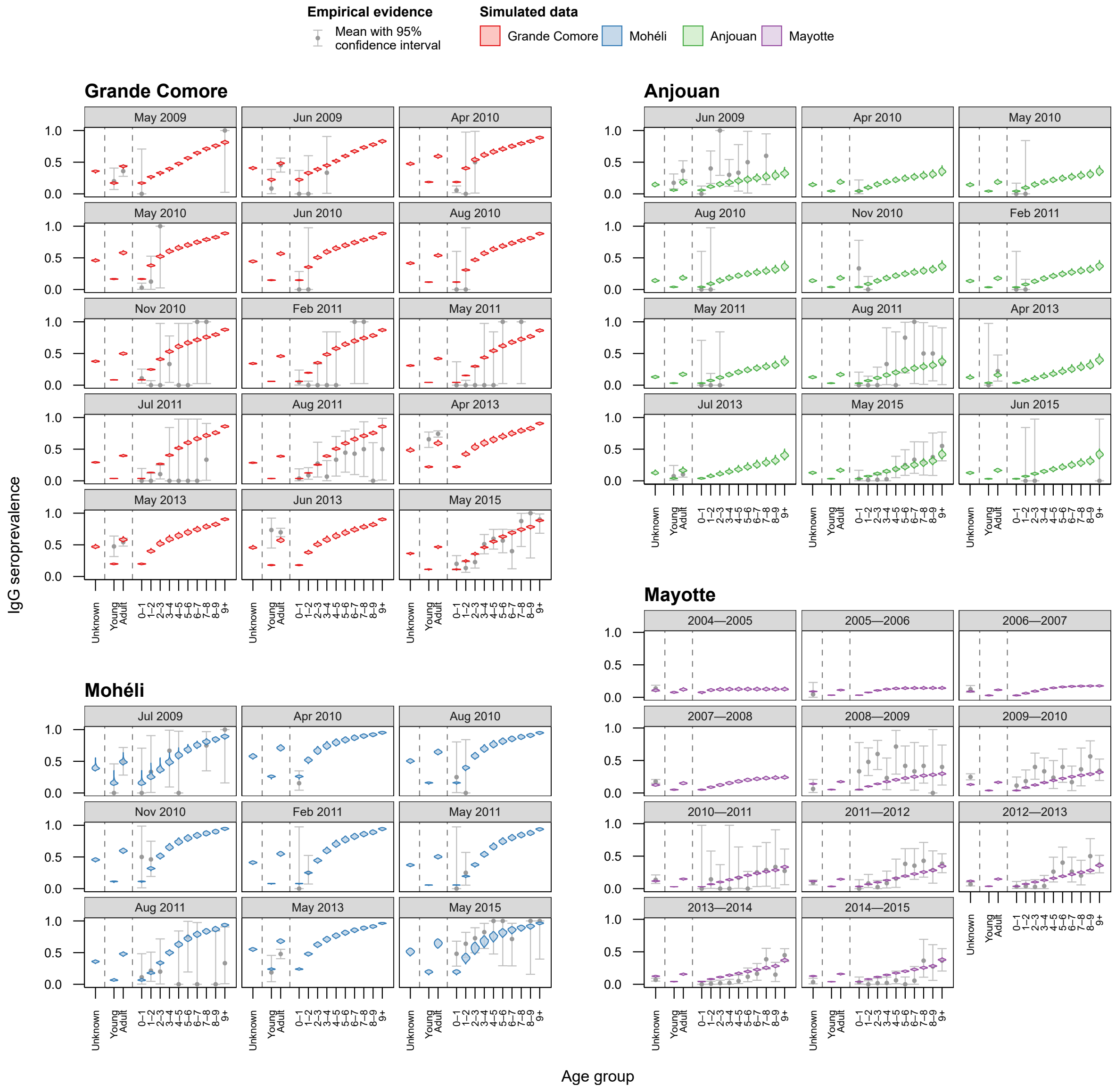


**Supplementary Figure 2: Model fit of the linear model (Model 2a) for each sero-survey conducted between July 2004 and June 2015.** The linear transmission model with the different seasonal components $\alpha$ and the same baseline transmission $\beta$ for each island fitted to the data third best out of all five tested models (DIC $= 1,433$). Shown is the fitted simulated seroprevalence for each aggregated sero-survey conducted throughout the study period (coloured violins). Seroprevalence of Grande Comore (red), Mohéli (blue) and Anjouan (green) were aggregated by month, and seroprevalence for Mayotte (purple) was aggregated by year. The black dots show the observed age-stratified IgG seroprevalence with 95% confidence interval (vertical bars). Simulated seroprevalence was generated through 1,000 realisations of the metapopulation model.

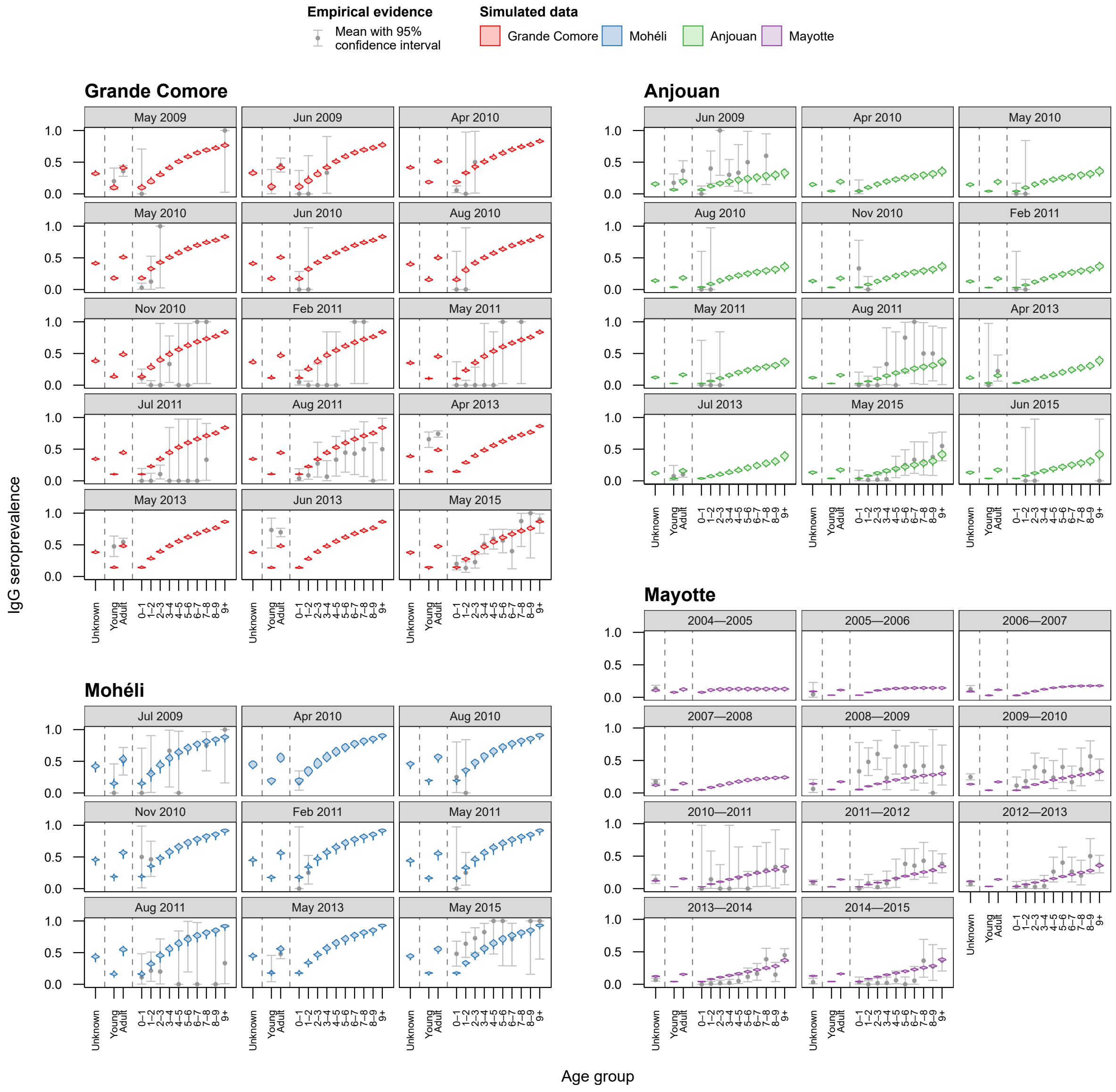


**Supplementary Figure 3: Model fit of the linear model (Model 2b) for each sero-survey conducted between July 2004 and June 2015.** The linear transmission model with the same seasonal component $\alpha$ and different baseline transmission $\beta$ for each island fitted to the data almost as poorly as the constant transmission model (DIC $= 1,578$). Shown is the fitted simulated seroprevalence for each aggregated sero-survey conducted throughout the study period (coloured violins). Seroprevalence of Grande Comore (red), Mohéli (blue) and Anjouan (green) were aggregated by month, and seroprevalence for Mayotte (purple) was aggregated by year. The black dots show the observed age-stratified IgG seroprevalence with 95% confidence interval (vertical bars). Simulated seroprevalence was generated through 1,000 realisations of the metapopulation model.

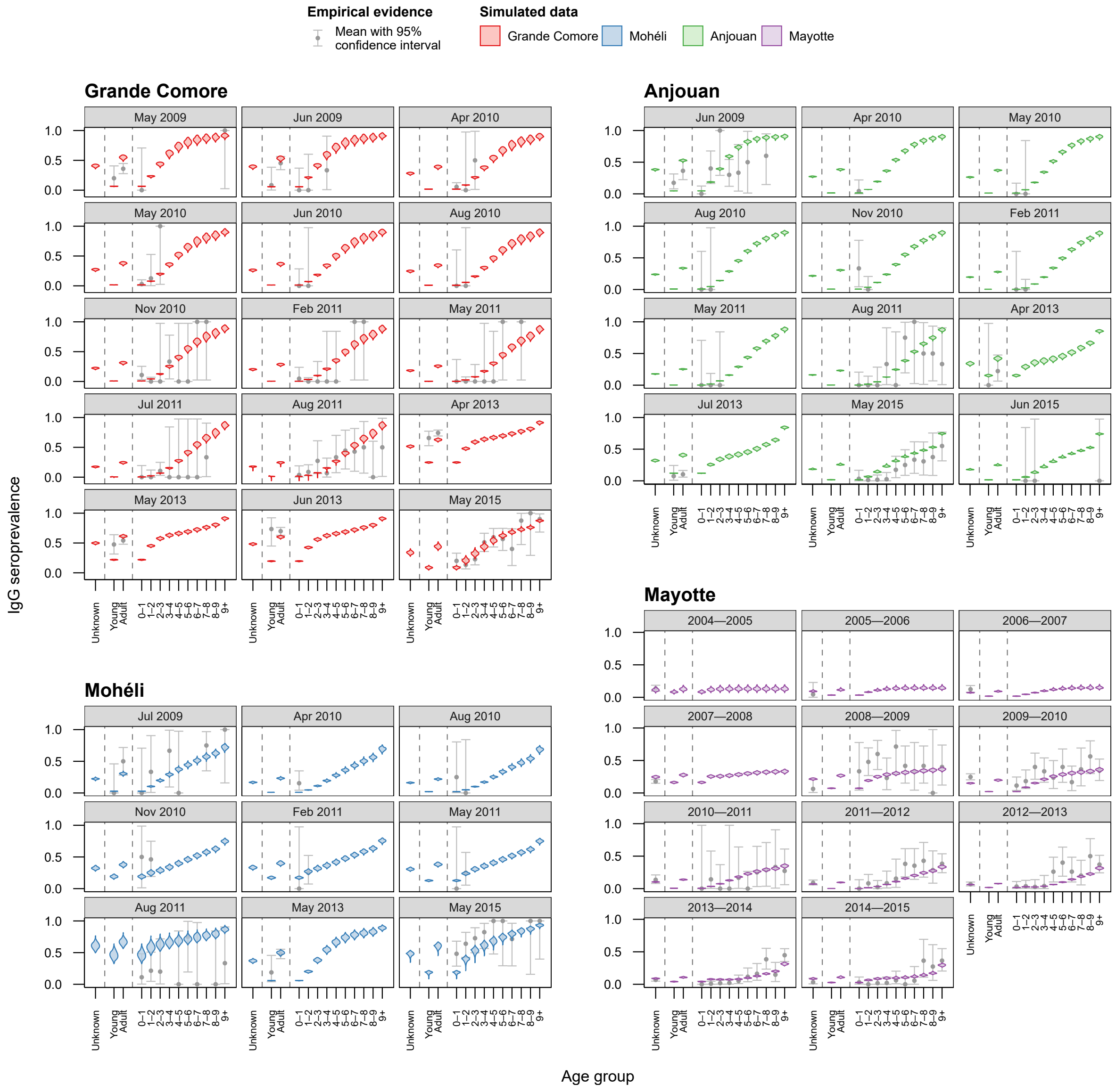


**Supplementary Figure 4: Model fit of the exponential model (Model 3a) for each sero-survey conducted between July 2004 and June 2015.** The exponential transmission model with the same seasonal component $\alpha$ and different baseline transmission $\beta$ for each island fitted to the data second best out of all five models tested (DIC $= 1,386$). Shown is the fitted simulated seroprevalence for each aggregated sero-survey conducted throughout the study period (coloured violins). Seroprevalence of Grande Comore (red), Mohéli (blue) and Anjouan (green) were aggregated by month, and seroprevalence for Mayotte (purple) was aggregated by year. The black dots show the observed age-stratified IgG seroprevalence with 95% confidence interval (vertical bars). Simulated seroprevalence was generated through 1,000 realisations of the metapopulation model.

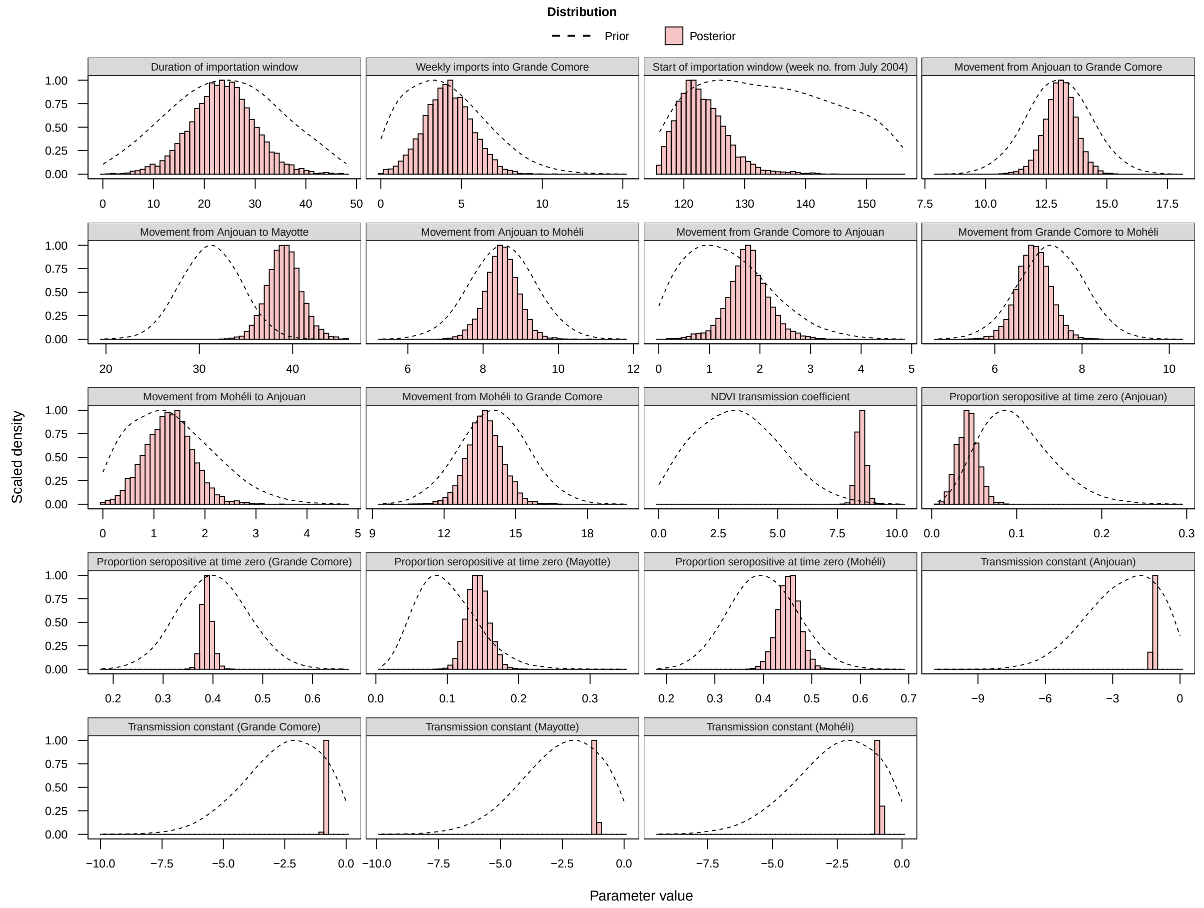


**Supplementary Figure 5: Posterior of estimated parameters in exponential model (Model 3b).** The exponential transmission model with the same seasonal component $a$ and different baseline transmission $b$ for each island fitted to the data best out of all five models tested. Shown are the prior (lines) and posterior (bars) distributions of each parameter. Prior distributions were informed by historical and current understanding of RVF epidemiology and livestock demography in the Comoros. Posterior distributions were generated from 10,000 samples from the fitted model.

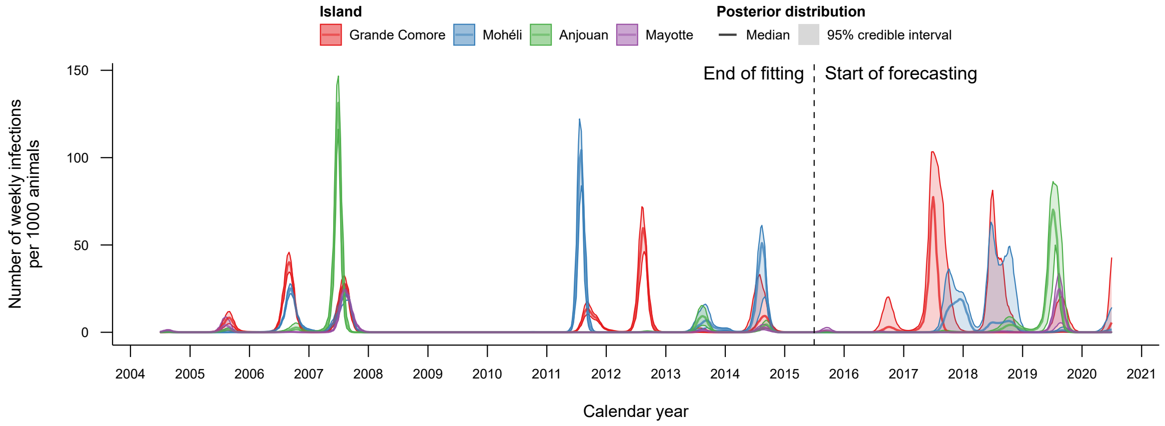


**Supplementary Figure 6: Weekly infections in the Comoros archipelago from 2004 to 2020.** The total number of weekly livestock infections per 1000 animals on each island from July 2004 to July 2020 as predicted by the exponential model (Model 3b). The estimated total livestock population on Grande Comore (red), Mohéli (blue), Anjouan (green) and Mayotte (purple) were 224,353; 31,872; 93,616 and 20,052 respectively. The median and 95% credible interval of infections was generated through 1,000 realisations of the metapopulation model.

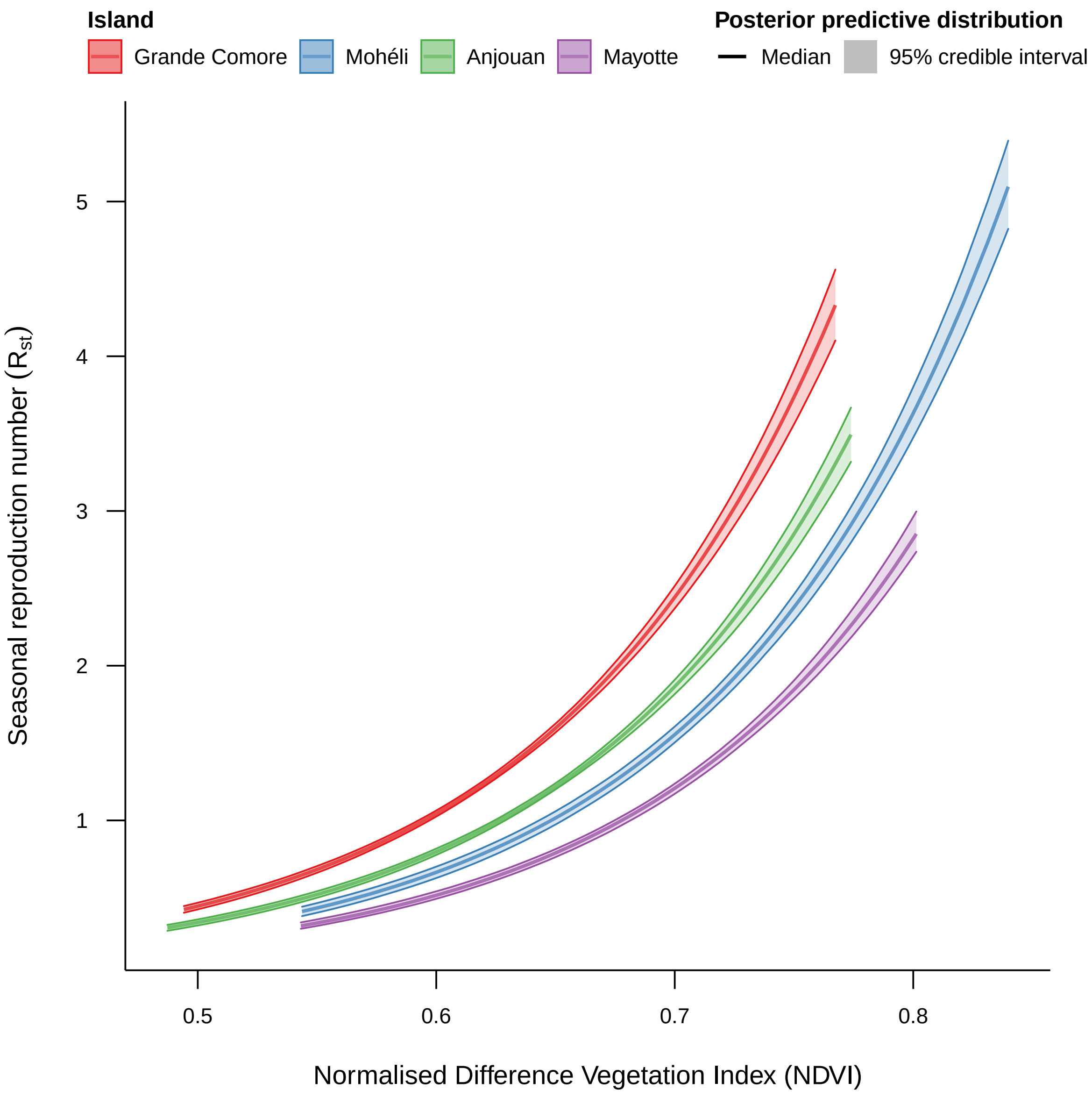


**Supplementary Figure 7: Relationship between NDVI and the seasonal reproduction number in the best fitted model (Model 3b).** In the best fitted model, within-island transmission was modelled as an exponential relationship to Normalized Difference Vegetation Index (NDVI). As a consequence, higher NDVI values resulted in higher seasonal reproduction numbers for the disease on Grande Comore (red), Mohéli (blue), Anjouan (green) and Mayotte (purple). The median and 95% credible interval of infections was generated through 1,000 realisations of the metapopulation model.

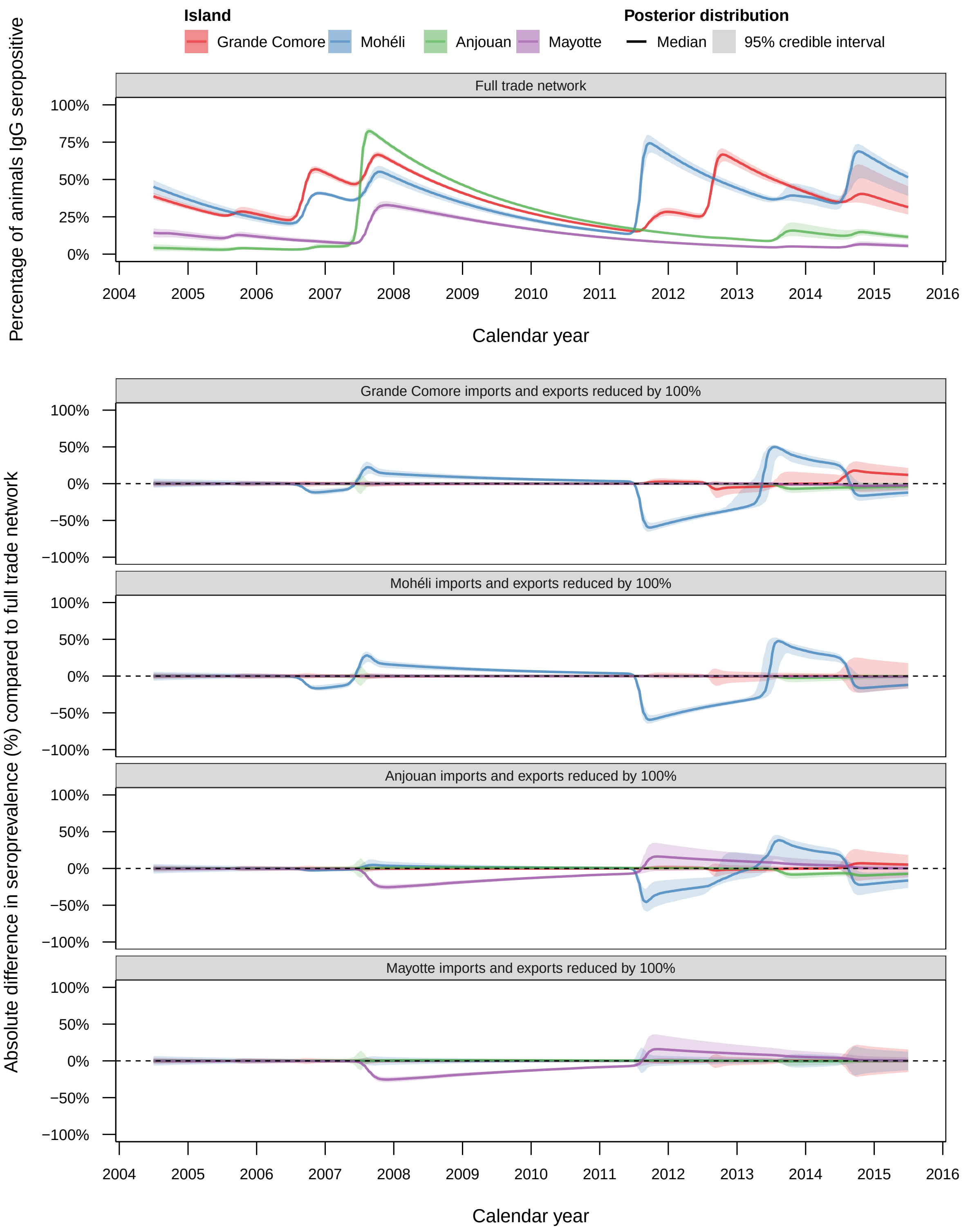


**Supplementary Figure 8: Effect of reducing imports and exports of each island on seroprevalence from 2004–2015.** The best fitted model (Model 3b) was simulated with different levels of import and export restrictions placed on each island. Shown is the absolute difference in the percentage of animals IgG seropositive for RVF from 2004–2015 under each movement restriction scenario on Grande Comore (red), Mohéli (blue), Anjouan (green) and Mayotte (purple) compared with the full trade network model. Distributions were generated from 1,000 realisations of each scenario.

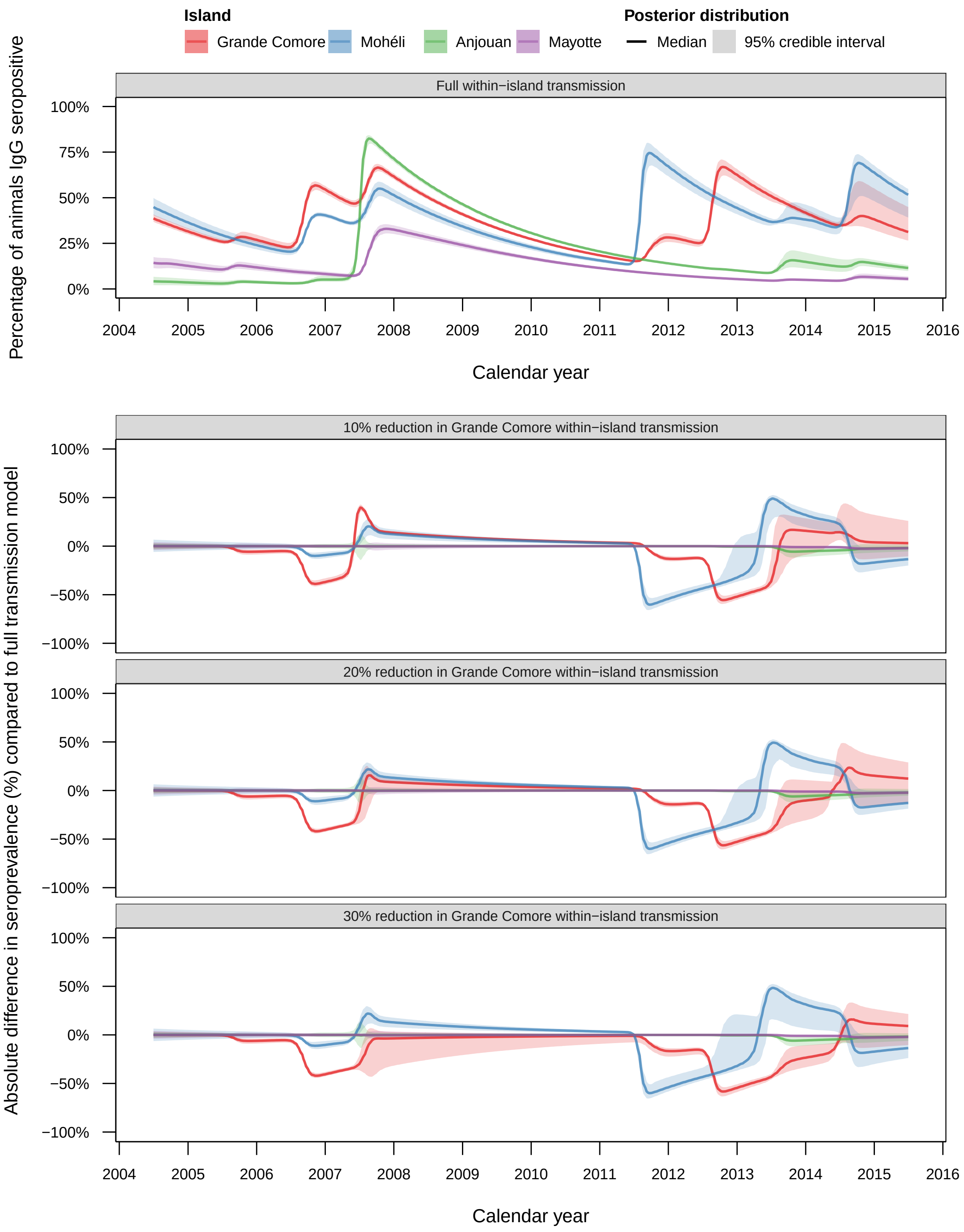


**Supplementary Figure 9: Effect of reducing within-island transmission of Grande Comore on seroprevalence from 2004–2015.** The best fitted model (Model 3b) was simulated with different reductions in the transmission rate on Grande Comore: 10%, 20% and 30%. Shown is the absolute difference in the percentage of animals IgG seropositive for RVF from 2004–2015 under each control scenario on Grande Comore (red), Mohéli (blue), Anjouan (green) and Mayotte (purple) compared with the full transmission model. Distributions were generated from 1,000 realisations of each scenario.

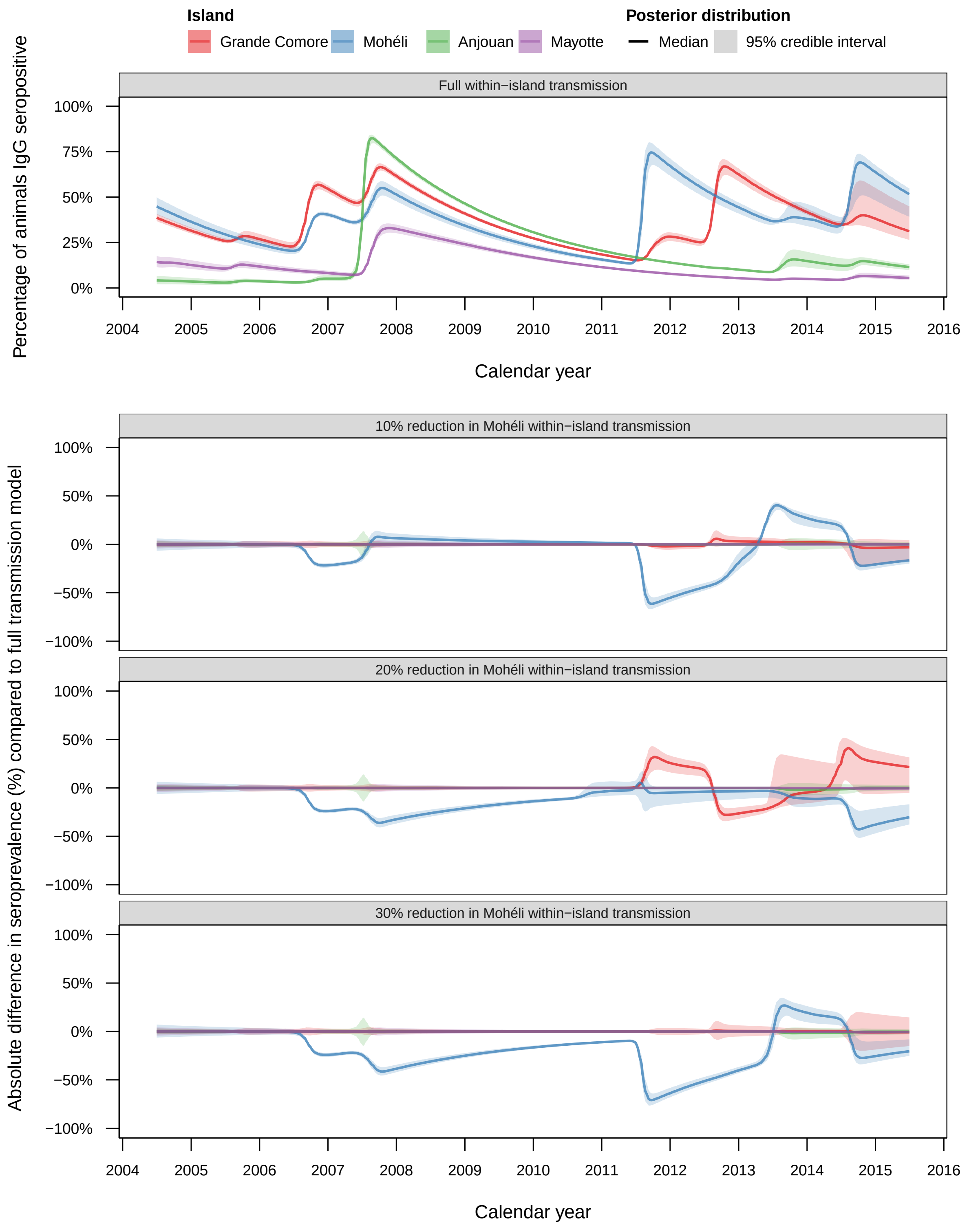


**Supplementary Figure 10: Effect of reducing within-island transmission of Mohéli on seroprevalence from 2004–2015.** The best fitted model (Model 3b) was simulated with different reductions in the transmission rate on Mohéli: 10%, 20% and 30%. Shown is the absolute difference in the percentage of animals IgG seropositive for RVF from 2004–2015 under each control scenario on Grande Comore (red), Mohéli (blue), Anjouan (green) and Mayotte (purple) compared with the full transmission model. Distributions were generated from 1,000 realisations of each scenario.

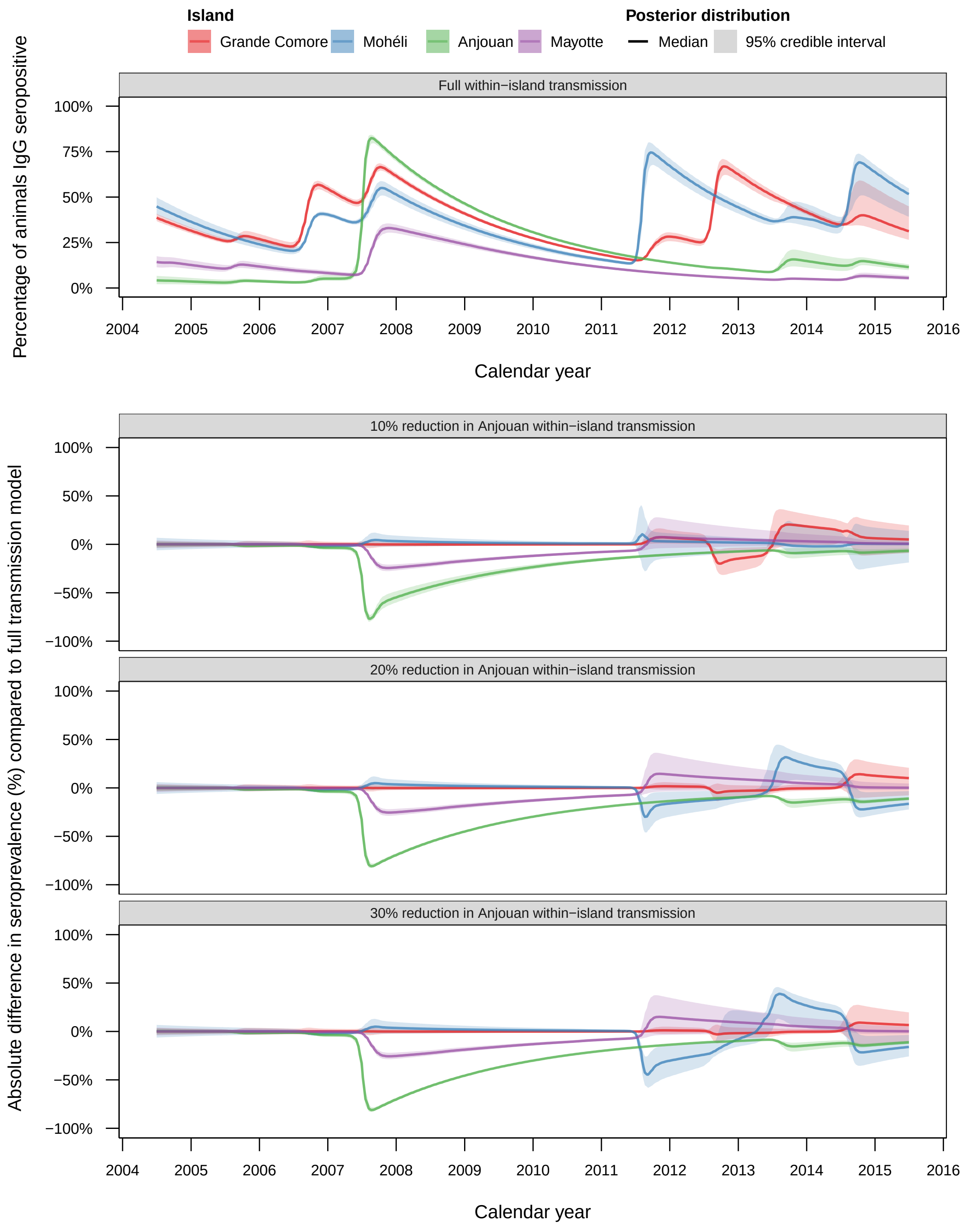


**Supplementary Figure 11: Effect of reducing within-island transmission of Anjouan on seroprevalence from 2004–2015.** The best fitted model (Model 3b) was simulated with different reductions in the transmission rate on Anjouan: 10%, 20% and 30%. Shown is the absolute difference in the percentage of animals IgG seropositive for RVF from 2004–2015 under each control scenario on Grande Comore (red), Mohéli (blue), Anjouan (green) and Mayotte (purple) compared with the full transmission model. Distributions were generated from 1,000 realisations of each scenario.

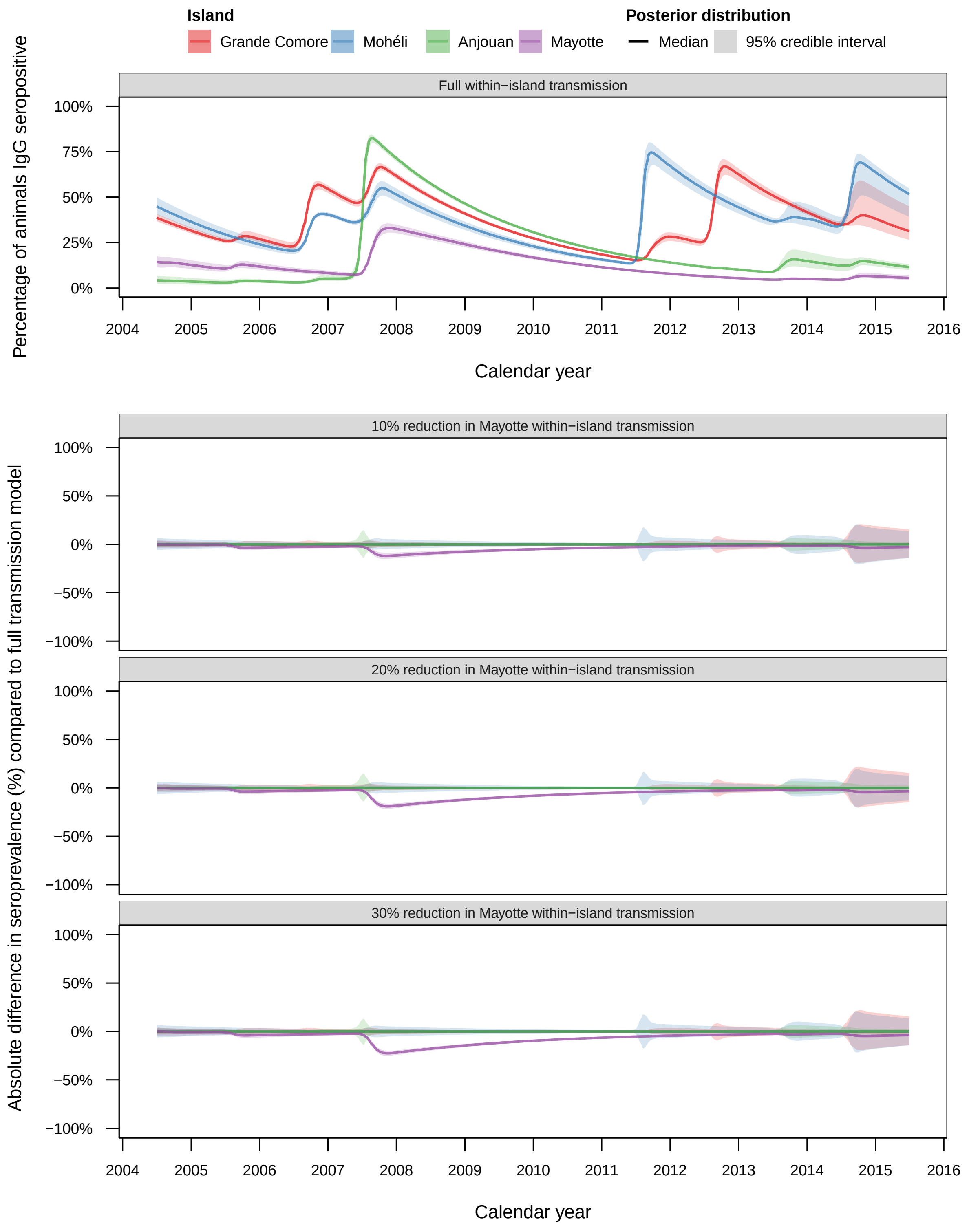


**Supplementary Figure 12: Effect of reducing within-island transmission of Mayotte on seroprevalence from 2004–2015.** The best fitted model (Model 3b) was simulated with different reductions in the transmission rate on Mayotte: 10%, 20% and 30%. Shown is the absolute difference in the percentage of animals IgG seropositive for RVF from 2004–2015 under each control scenario on Grande Comore (red), Mohéli (blue), Anjouan (green) and Mayotte (purple) compared with the full transmission model. Distributions were generated from 1,000 realisations of each scenario.
